# Two parallel sRNA amplification cycles contribute to RNAi inheritance in *C. elegans*

**DOI:** 10.1101/2021.08.13.456232

**Authors:** John Paul Tsu Ouyang, Wenyan Zhang, Geraldine Seydoux

**Affiliations:** HHMI and Dept. of Molecular Biology and Genetics, Johns Hopkins University School of Medicine, Baltimore MD USA

**Keywords:** RNA-mediated interference, RNAi inheritance, small RNA amplification, Argonautes, nuage, helicase

## Abstract

RNA-mediated interference (RNAi) is a conserved mechanism that uses small RNAs (sRNAs) to tune gene expression. In *C. elegans*, exposure to dsRNA induces the production of gene-specific sRNAs that are propagated to progeny not exposed to the dsRNA trigger. We present evidence that RNAi inheritance is mediated by two parallel sRNA amplification loops. The first loop, dependent on the nuclear Argonaute HRDE-1, targets nascent transcripts, and reduces but does not eliminate productive transcription at the locus. The second loop, dependent on the conserved helicase ZNFX-1, targets mature transcripts and concentrates them in perinuclear condensates (nuage). Each amplification loop generates a distinct class of sRNAs, with the ZNFX-1 loop responsible for the bulk of sRNA production on the region targeted by the trigger. By independently targeting nascent and mature transcripts, the HRDE-1 and ZNFX-1 loops ensure maximum silencing in progeny not exposed to the trigger.

## Introduction

RNA-mediated interference (RNAi) is a wide-spread mechanism that employs small RNAs (sRNAs) to modulate gene expression. First discovered as a response to exogenously provided dsRNA in *C. elegans* (Fire et al., 1998), RNAi-like mechanisms have been described in a broad range of organisms, from yeast to mammals. At the core of the RNAi machinery are RNA-induced silencing complexes (RISC) consisting of ∼20-base single stranded RNAs complexed with Argonaute proteins. RISC recognizes complementary RNAs via base pairing with the sRNA guide and effects silencing by reducing RNA stability and/or translation efficiency (Bartel, 2018; Billi et al., 2014; Carthew & Sontheimer, 2009). Certain RISC complexes also recognize nascent transcripts and interfere with productive transcription by stalling RNA polymerase, RNA processing and/or recruiting chromatin modifiers to the locus (Billi et al., 2014; Castel & Martienssen, 2013; Weiser & Kim, 2019). In many organisms, sRNA pathways depend on cycles that amplify the production of sRNAs to achieve maximal silencing (Billi et al., 2014; Czech & Hannon, 2016). In *Drosophila*, a complex “ping pong” cycle in perinuclear condensates amplifies the processing of genomically-encoded precursor transcripts containing sRNAs that target active transposable elements (piRNAs; Czech and Hannon, 2016). In *S. pombe*, the nuclear RISC-like complex (RITS) recruits an RNA-dependent RNA polymerase (RdRP) to the targeted locus (Martienssen & Moazed, 2015). The RdRP uses nascent transcripts as templates for continued synthesis of sRNAs that feed back into RITS (Martienssen & Moazed, 2015). In both cases, the sRNA amplification loops depend on transcription of the locus targeted for silencing to supply the template necessary to stimulate the processing (*Drosophila*) or the *de novo* synthesis (*S. pombe*) of the relevant sRNAs.

As in *S. pombe*, sRNA amplification in *C. elegans* involves the activity of RdRPs that synthesize new sRNAs on transcripts recognized by RISC complexes. Two sRNA amplification mechanisms have been described. A first mechanism involves “primary” sRNAs derived from genomically encoded loci (e.g. piRNAs) or from dsRNA processed by the RNA endonuclease Dicer (Billi et al., 2014). Recognition of complementary transcripts by primary sRNAs, complexed with primary Argonautes (e.g. RDE-1), leads to their cleavage by the endonuclease RDE-8 and tailing of the 5’ fragment by the poly(UG) polymerase MUT-2/RDE-3 (Shukla et al., 2020; Tsai et al., 2015). The “pUG” tail recruits RdRPs that synthesize “secondary” sRNAs near the cleavage site (Shukla et al., 2020). Secondary sRNAs in turn associate with secondary Argonautes (WAGOs) to trigger the degradation of complementary transcripts in the cytoplasm by an unknown mechanism (Yigit et al., 2006). A second cycle depends on the Argonaute HRDE-1, which shuttles between the cytoplasm and nucleus (Ashe et al., 2012; Buckley et al., 2012; Shirayama et al., 2012; Sapetschnig et al., 2015). This cycle is less well understood but is thought to function similarly to RITS in *S. pombe*, coordinating heterochromatin deposition and sRNA synthesis by binding to nascent transcripts (Billi et al., 2014; Martienssen & Moazed, 2015; Weiser and Kim, 2019).

A fascinating aspect of RNAi-induced silencing in *C. elegans* is the ability for the silenced state to be passed on to progeny even in the absence of the initial trigger (Fire et al., 1998; Grishok et al., 2000; Alcazar et al., 2008; Lev et al., 2017; Weiser and Kim, 2019). pUGylated transcripts have been observed in the progeny of worms exposed to dsRNA, raising the possibility that a pUGylation-dependent sRNA amplification cycle may be heritable (Shukla et al., 2020, 2021). An early study examining RNAi in somatic tissues of *C. elegans* suggested, however, that only primary Argonautes can initiate sRNA amplification (Pak et al., 2012); secondary Argonautes, in contrast, only target cognate mRNAs for degradation (Pak et al., 2012). Subsequent studies showed that production of “tertiary” sRNAs is allowed in the germline and depends on HRDE-1 and other nuclear RNAi factors (Sapetschnig et al., 2015). Unlike secondary sRNAs which map near the site of the primary sRNA trigger, tertiary sRNAs generated through the nuclear RNAi pathway map throughout the transcript possibly because they are synthesized by a nuclear RdRP that uses nascent transcripts as templates (Sapetschnig et al., 2015). Cytoplasmic factors have also been implicated in RNAi inheritance including the Argonaute WAGO-4 and the helicase ZNFX-1 (Ishidate et al., 2018; Wan et al., 2018; Xu et al., 2018). Whether these factors function in the HRDE-1 cycle or a different sRNA amplification cycle was not known.

In germ cells, several components of the RNAi machinery are concentrated in condensates at the nuclear periphery. Five condensate types have been described so far, including P granules (Strome & Wood, 1982), Mutator foci (Phillips et al., 2012), R2 bodies (Yang et al., 2014), Z granules (Wan et al., 2018), and SIMR foci (Manage et al., 2020). In this study, we refer to these condensates collectively as “nuage” following the convention for perinuclear condensates in *Drosophila* and mammalian systems (Dodson and Kennedy, 2020). Nuage condensates often overlay nucleopores and have been proposed to serve as compartments specialized in transcript surveillance and sRNA amplification (Voronina et al., 2011; Gao and Arkov, 2013; Dodson and Kennedy, 2020; Sundby et al., 2021). What specific functions these compartments play in RNAi inheritance and sRNA amplification is not known.

In this study, we examined the fate of germline mRNAs in animals exposed (by feeding) to a gene-specific dsRNA trigger. Our findings indicate that the HRDE-1 cycle, although sufficient to partially silence the locus, is not sufficient for robust inheritance of the silenced state. A second cycle involving the Z granule component ZNFX-1 is also required in parallel. We find that ZNFX-1 is responsible for localization of targeted mRNAs to nuage, and to maintain pUGylation and sRNA amplification in progeny. Together, our findings suggest a model where nuage condensates represent privileged compartments where non-primary sRNAs are permitted to initiate new rounds of pUGylation and sRNA amplification on mature transcripts exported from the nucleus.

## Results

### Changes in the abundance of nascent and cytoplasmic transcripts appear within hours of exposure to the dsRNA trigger

To examine the consequences of RNAi-induced gene silencing, we first used fluorescent *in situ* hybridization (FISH) to examine changes in the level and localization of a targeted transcript. We chose *mex-6* as a model transcript because 1) *mex-6* is expressed in the pachytene region of the adult germline, where nuage condensates are prominent (Fig 1), 2) *mex-6* is minimally targeted by endogenous sRNAs under non-RNAi conditions (Fig S1A), and 3) *mex-6* is a non-essential maternal-effect gene (redundant with *mex-5*), whose silencing does not affect germline development or morphology (Schubert et al., 2000).

**Figure 1.**
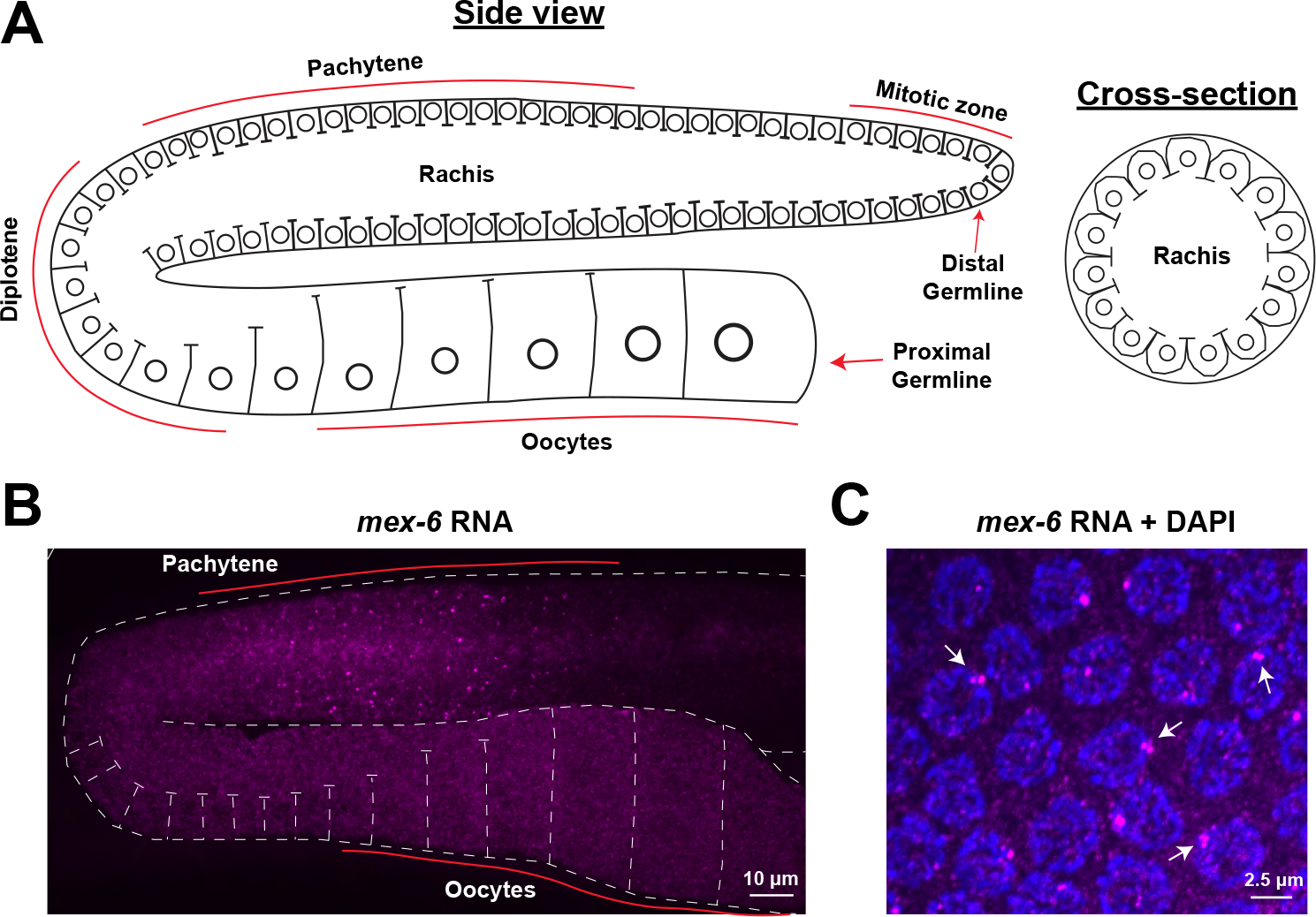
*mex-6* RNA expression in the C. elegans germline. A) Schematic diagrams (adapted from Gartner et al., 2008) of adult hermaphrodite gonads from a side or cross-sectional view. Circles indicate germline nuclei; lines indicate plasma membranes. Distal nuclei are in mitosis and progress through the different stages of meiosis and oogenesis as they move towards the proximal end of the germline. A common cytoplasm (rachis) runs through the entire germline, excluding the most distal (mature) oocyte which is completely cellularized. B) Maximum projection photomicrograph of an adult *C. elegans* germline oriented as in A and with *mex-6* RNA visualized by fluorescent in situ hybridization (magenta). C) High resolution photomicrograph showing pachytene nuclei (blue, stained with DAPI) and *mex-6* RNA (magenta). Arrows point to nuclei where nascent transcripts at the *mex-6* locus resolve into two or more closely apposed foci, as expected for tightly synapsed, replicated homologous chromosomes in the pachytene stage of meiosis.

The germline of adult *C. elegans* hermaphrodites is a syncytial tissue organized in two tubes folded into distal and proximal arms. Germ cells are arranged in order of maturation with mitotic stem cells at the distal-most end, followed by germ cells that have initiated meiosis (pachytene), and growing oocytes in the proximal arm (Fig. 1A). Like other maternal transcripts, *mex-6* RNA is transcribed in nuclei in the pachytene region and accumulates in the cytoplasm shared by pachytene nuclei (rachis of the distal arm) and growing oocytes (Chi and Reinke, 2006). As expected, by FISH, we detected *mex-6* transcripts diffuse in the rachis and cytoplasm of growing oocytes and concentrated in bright nuclear puncta within pachytene nuclei (but not in oocyte nuclei; Fig 1B, 1C). At high magnification, the nuclear puncta overlapped with DAPI staining and occasionally resolved into twin or triplet dots (Fig. 1C), consistent with tight pairing of replicated homologous chromosomes in pachytene nuclei (Lui & Colaiácovo, 2013). Two-color RNA FISH against *mex-6* and a transcript expressed from a linked locus (*puf-5*) revealed closely linked puncta in pachytene nuclei (Fig S1B), confirming that the nuclear puncta correspond to nascent transcripts at the *mex-6* and *puf-5* loci.

To silence the *mex-6* gene, we designed a 600bp double-stranded RNA trigger targeting the 3’ most region of the *mex-6* transcript, including the 3’ UTR. The *mex-6* trigger was designed to avoid overlap with the FISH probes used to visualize the *mex-6* transcript (Fig S2A). Synchronized first-day adult hermaphrodites were plated onto bacteria expressing the *mex-6* trigger (*mex-6* RNAi) or a control vector trigger ( Control RNAi). Animals were collected for FISH after 2, 4, 6, 8, and 24 hours of feeding and nuclear and cytoplasmic *mex-6* FISH signals were quantified (Methods). We first detected a reduction in FISH signal in the cytoplasm of oocytes after 4 hours of treatment, culminating in > 90% decrease by 24 hours (Fig 2A, 2B), confirming the efficacy of our RNAi feeding protocol. Starting at the 6-hour time point, we also detected an increase in the intensity distribution of nuclear puncta in the pachytene region (Fig 2A, 2C, 2D). The intensity distribution remained elevated through the 8-hour time point (Fig 2C, 2D), before diminishing slightly by the 24-hour time-point (Fig 2A, 2C). We conclude that, in the first 24 hours of exposure to the dsRNA trigger, RNAi induces a transient increase in the accumulation of nascent transcripts at the locus and a steady decrease in cytoplasmic transcripts.

**Figure 2.**
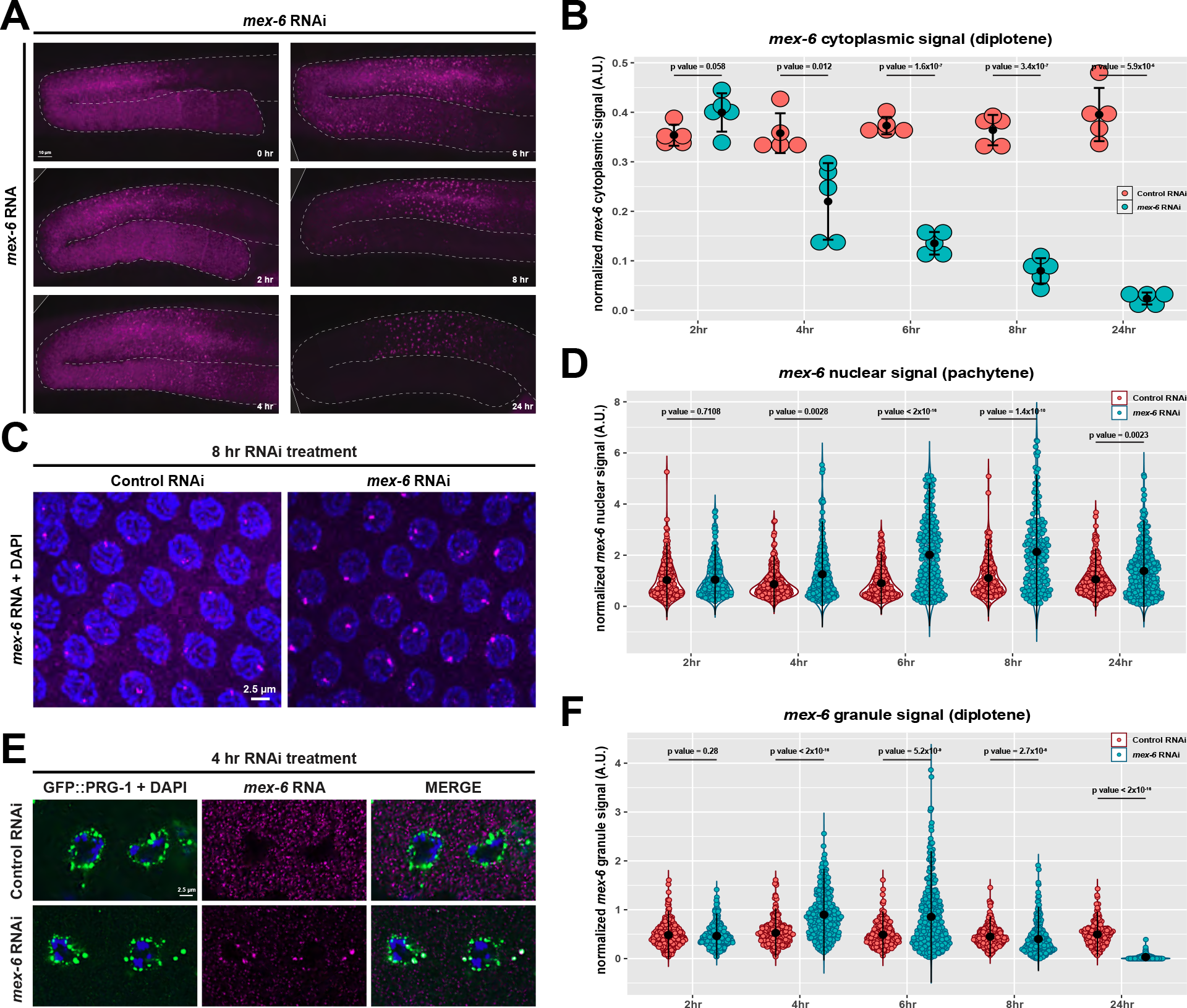
Evolution of *mex-6* RNA during a 24-hour RNAi treatment in wild-type hermaphrodites (P0 generation). A) Maximum projection photomicrograph of germlines oriented as in Figure 1 with *mex*-6 RNA (magenta) at indicated times post onset of feeding RNAi. B) Graph comparing mean cytoplasmic *mex*-6 RNA FISH signals (diplotene region) in control (red) and *mex*-6 (blue) RNAi conditions at the indicated time points of RNAi treatment. Each dot represents a single germline. Values (arbitrary units) were normalized to *puf*-5 RNA FISH signals visualized in same germline (Methods). Central black dot and error bars represent the mean and standard deviation, respectively. P values were calculated using an unpaired t-test. C) Maximum projection photomicrographs of pachytene nuclei showing *mex*-6 RNA (magenta) and DNA (blue, stained with DAPI) after 8 hours of RNAi treatment. D) Graph comparing maximum nuclear *mex*-6 RNA FISH signals (pachytene region) in control (red) vs *mex*-6 (blue) RNAi conditions at the indicated time points of RNAi treatment. Each dot represents one nucleus. Nuclei in three worms were quantified for each time point and condition. Values (arbitrary units) were normalized to *puf*-5 RNA FISH signals visualized in same nuclei (Methods). Central black dot and error bars represent the mean and standard deviation respectively. P values were calculated using an unpaired Wilcoxon test. E) Photomicrographs of two oocytes with the germ granule marker GFP::PRG-1 in green, DNA in blue (DAPI) and *mex*-6 RNA in magenta after 4 hours of RNAi treatment. F) Graph comparing average *mex*-6 RNA FISH signal in germ granules in control (red) vs *mex*-6 (blue) RNAi conditions at the indicated time points of RNAi treatment. Each dot corresponds to a granule. Five worms were quantified for each stage and condition. Values (arbitrary units) were normalized to *puf*-5 RNA FISH signals visualized in same granules (Methods). Central black dot and error bars represent the mean and standard deviation, respectively. P values were calculated using an unpaired Wilcoxon test.

### Targeted transcripts accumulate in nuage

Beginning at the 4-hour time point, we also noticed accumulation of *mex-6* transcripts in micron-sized clusters in the cytoplasm of growing oocytes (Fig 2A, 2E, 2F). Colocalization with nuage markers ZNFX-1 and PRG-1 revealed that the clusters coincide with nuage (Fig 2E, S2B). At the 6 and 8-hour time points, we also detected *mex-6* accumulation in nuage in the loop and pachytene region (Fig 2A). At the 24-hour time point, *mex-6* accumulation in nuage in oocytes was strongly diminished, mirroring the strong depletion of *mex-*6 transcripts in the cytoplasm (Fig 2A, 2B). However, *mex-6* signal could still be detected in nuage in the pachytene region where the *mex-6* locus is transcribed (Fig S2C). We conclude that *mex-6* transcripts accumulate in nuage throughout the RNAi response. Note that the resolution of our *in situ* protocol did not allow us to distinguish whether targeted RNAs localized to all or a subset of condensate types in nuage.

### RNAi-induced changes in nascent and cytoplasmic transcripts require *rde-1* and *mut-16* activity

To determine whether the changes observed were dependent on the RNAi machinery, we examined *rde-1* and *mut-16* mutants. RDE-1 is the Argonaute that recognize primary sRNAs derived from exogenous triggers (Tabara et al., 1999; Yigit et al., 2006), and MUT-16 is required for amplification of secondary sRNAs (Zhang et al., 2011). We found that *rde-1* and *mut-16* mutants were completely defective in the RNAi response (Fig S2D, S2E). *rde-1* and *mut-16* animals exposed to the *mex-6* trigger resembled animals exposed to the no-RNAi control trigger at both the 8 and 24-hour time points and showed none of the changes observed in wild-type animals exposed to the dsRNA trigger (Fig S2D, S2E). We conclude that changes in the level and localization of nascent and cytoplasmic transcripts depend on initiation of the RNAi response by RDE-1 and synthesis of secondary sRNAs.

### RNAi-induced changes in nascent and cytoplasmic transcripts are inherited in F1 and F2 generations

To examine inheritance of the RNAi response, we performed FISH analysis on the progeny (F1) of hermaphrodites fed the dsRNA trigger (P0). P0 hermaphrodites at the 24-hour time point were bleached, and F1 embryos were synchronized and plated onto non-RNAi plates starting at the L1 stage. The F1s were raised to the adult stage (in the absence of the RNAi trigger) and processed for FISH. We observed a strong reduction in *mex-6* RNA throughout the germline of F1 animals compared to F1 progeny of animals exposed to control RNAi (Fig. 3A, S3A). The average intensity of nuclear signals in the pachytene region was reduced by ∼50% compared to controls (Fig 3B). Despite this strong reduction, we still detected transcripts in perinuclear dots overlapping with nuage markers in the pachytene region (Fig 3C, S3B). In contrast, little to no nuage accumulation was evident in oocytes (Fig 3A, S3C). Similar observations were made in the F2 generation (Fig S3C). These observations suggest that, despite a reduction in nascent transcripts, some *mex-6* transcripts are still exported from the nucleus and allowed to accumulate at least transiently in nuage in the pachytene region in F1 and F2 animals.

**Figure 3.**
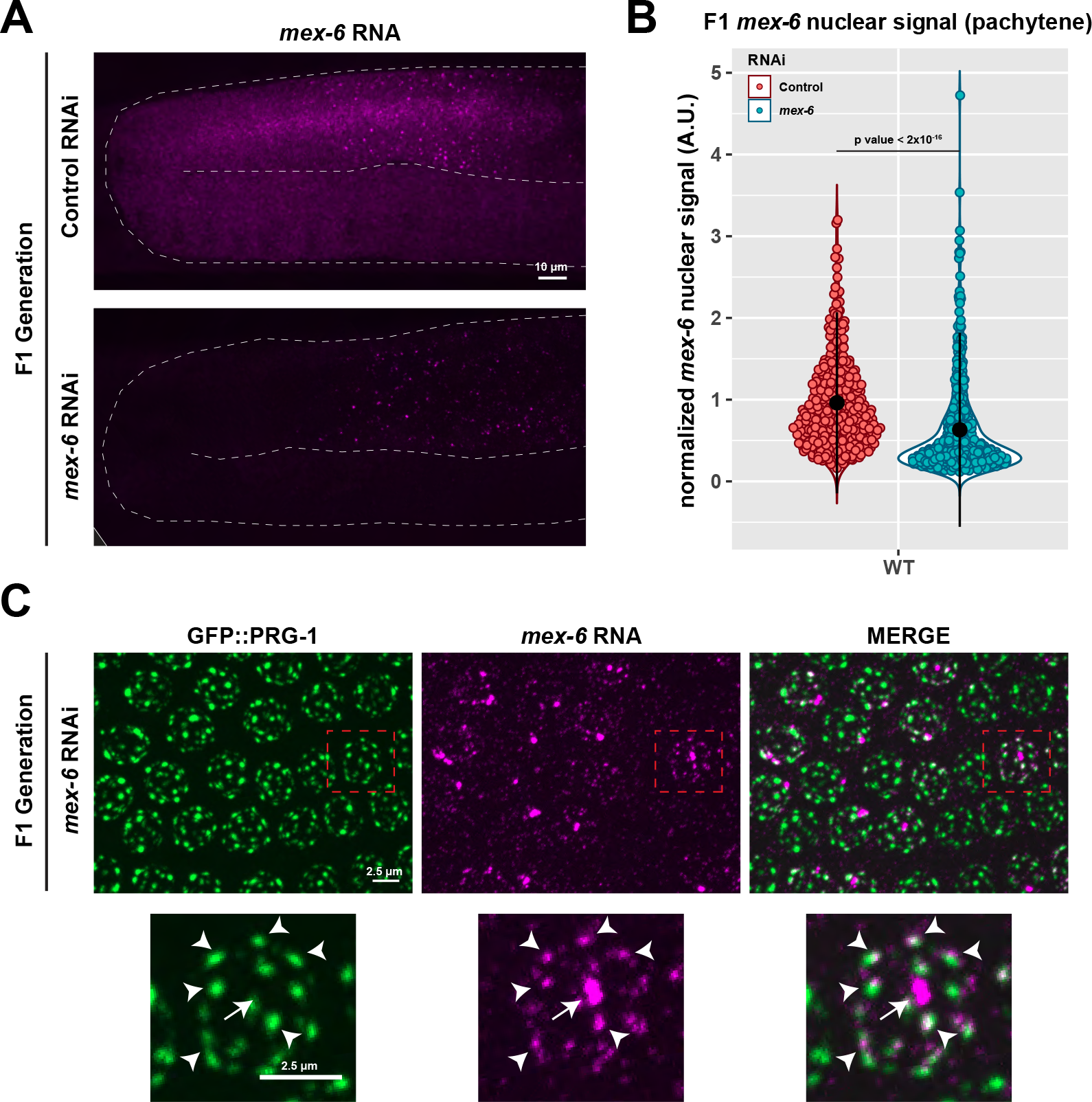
*mex-6* RNA in adult progeny (F1 generation) of animals exposed to *mex-6* dsRNA. A) Maximum projection photomicrographs of germlines showing *mex-6* RNA (magenta) in adult F1 progeny of animals exposed to control or *mex-6* RNAi (Methods). B) Graph comparing maximum nuclear *mex-6* RNA FISH signals (pachytene region) in F1 progeny of animals exposed to control (red) or *mex-6* (blue) RNAi. Each dot represents one nucleus. Nuclei in three worms were quantified for each time point and condition. Values (arbitrary units) were normalized to *puf-5* RNA FISH signals visualized in same nuclei (Methods). Central black dot and error bars represent the mean and standard deviation respectively. P values were calculated using an unpaired Wilcoxon test. C) Maximum projection photomicrographs of pachytene nuclei showing the germ granule marker GFP::PRG-1 (green) and *mex-6* RNA (magenta) in F1 progeny of animals exposed to *mex-6* RNAi. See Fig. S3B for photomi- crographs of the control condition. Arrow points to *mex-6* RNA signal at the locus and arrow heads point to *mex-6* RNA foci overlapping with perinuclear nuage. Note that the *mex-6* RNA signal overlaps but is not perfectly coincident with PRG-1, a P granule marker, suggesting that *mex-6* RNA may reside in a different nuage compartment.

### *hrde-1* is required for nascent transcripts to respond to RNAi in P0 and F1 animals

HRDE-1/WAGO-9 is a germline-specific nuclear Argonaute required for inheritance of the RNAi-induced silenced state (Ashe et al., 2012; Buckley et al., 2012; Shirayama et al., 2012). We found that *hrde-1* mutants mounted a normal RNAi response in oocytes of P0 hermaphrodites: we observed loss of *mex-6* RNA in the cytoplasm and accumulation in nuage in *hrde-1* mutants as in wild-type (Fig 4A, S4A). We observed, however, no change in the intensity distribution of nuclear puncta in the pachytene region (Fig 4A, 4B, 4C), consistent with a failure to silence the *mex-6* locus. To explore this possibility further, we examined the accumulation of *mex-6* transcripts in the rachis, the shared cytoplasm immediately adjacent to pachytene nuclei. In wild-type animals, *mex-6* levels in the rachis declined by >90% at the 24-hour time point (Fig 4D, 4E). In contrast, in *hrde-1* mutants, *mex-6* levels in the rachis were reduced only by ∼50% at the 24-hour time point (Fig 4D, 4E). These observations suggest that *hrde-1* mutants fail to interfere with the production of *mex-6* transcripts in P0 hermaphrodites. We obtained similar results in a strain mutated for another component of the nuclear RNAi machinery, *nrde-2* (Fig S4B).

**Figure 4.**
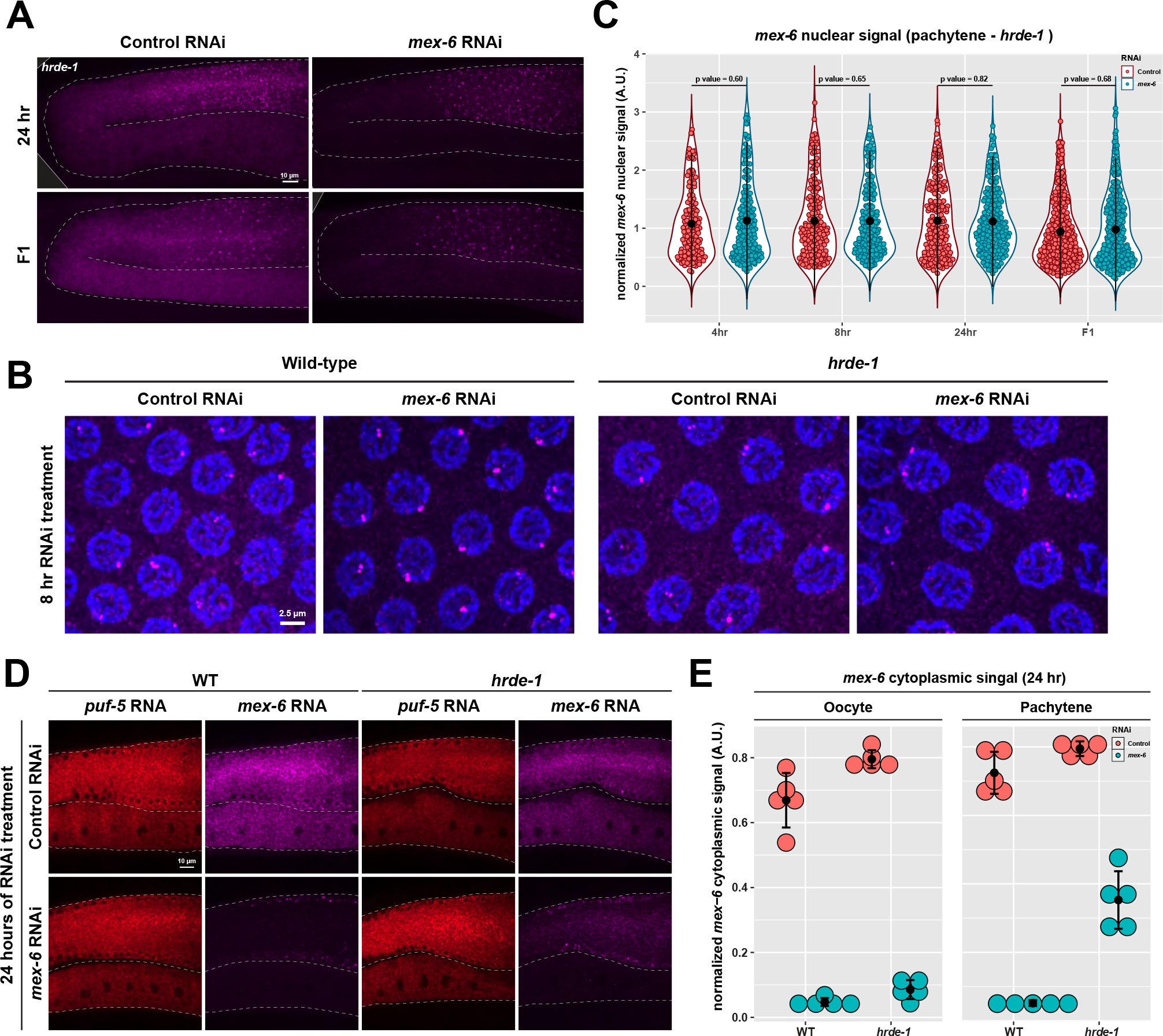
*hrde-1* is required for RNAi-induced changes in nascent transcripts. A) Maximum projection photomicrographs of germlines showing *mex-6* RNA (magenta) in P0 (24 hr RNAi exposure) and F1 *hrde-1* mutants under control or *mex-6* RNAi conditions. B) Maximum projection photomicrographs of pachytene nuclei in P0 wild-type and *hrde-1* mutant animals showing *mex-6* RNA (magenta) and DNA (blue, stained with DAPI) following 8 hours of either control or *mex-6* RNAi treatment. C) Graph comparing maximum nuclear *mex-6* RNA FISH signals (pachytene region) in P0 hrde-1 mutants following either control (red) or *mex-6* (blue) RNAi at the indicated timepoints. Each dot represents one nucleus. Three worms were quantified for each stage and condition. Central black dot and error bars represent the mean and standard deviation, respectively. Values (arbitrary units) were normalized to *puf-5* RNA FISH signals visualized in same nuclei (Methods). P values were calculated using an unpaired Wilcoxon test. D) Single z-plane photomicrographs showing *mex-6* RNA (magenta) and control *puf-5*RNA (red) in the cytoplasm in the pachytene and oocyte regions comparing wild-type and *hrde-1* mutants at 24 hours of RNAi treatment. *mex-6* RNA is depleted in both regions in wild-type but is only partially depleted in the pachytene region of *hrde-1* mutants consistent with a failure to silence the locus. E) Graph comparing mean *mex-6* RNA levels in the cytoplasm of oocytes and pachytene region in wild-type and *hrde-1* mutant at 24 hours of RNAi treatment. Each dot represents an individual animal. Values (arbitrary units) were normalized to *puf-5* RNA FISH signals visualized in the same areas (Methods). Central black dot and error bars represent the mean and standard deviation, respectively.

Failure to silence the *mex-6* locus was also observed in *hrde-1* F1 progeny. The distribution of nuclear puncta intensity was similar in *hrde-1* F1s and no-RNAi controls (Fig 4A, 4C). As in wild-type, however, *hrde-1* F1 progeny accumulated *mex-6* transcripts in nuage in the pachytene region (Fig S4C). *mex-6* RNA levels in the pachytene rachis were higher in *hrde-1* F1 progeny than in wild-type F1s but averaged only 50% of that observed in the no-RNAi controls (Fig S4D). We conclude that *hrde-1* is required for silencing of the locus in P0 and F1 animals (nuclear response) but is not essential for RNA degradation in the cytoplasm and accumulation in nuage in P0 and F1 animals (cytoplasmic response).

### *znfx-1* is required for accumulation of targeted transcripts in nuage in P0 and F1 animals

ZNFX-1 is an SF1 helicase-domain containing zinc finger protein that, like HRDE-1, is required for inheritance of the RNAi-induced silenced state (Ishidate et al., 2018; Wan et al., 2018). Unlike HRDE-1, which is primarily nuclear, ZNFX-1 localizes to specific condensates in nuage called Z granules (Wan et al., 2018). We found that, in *znfx-1* P0 animals, *mex-6* transcripts were rapidly degraded as in wild-type (Fig S5A). *mex-6* transcripts, however, failed to accumulate in nuage (Fig 5A, 5B).

**Figure 5.**
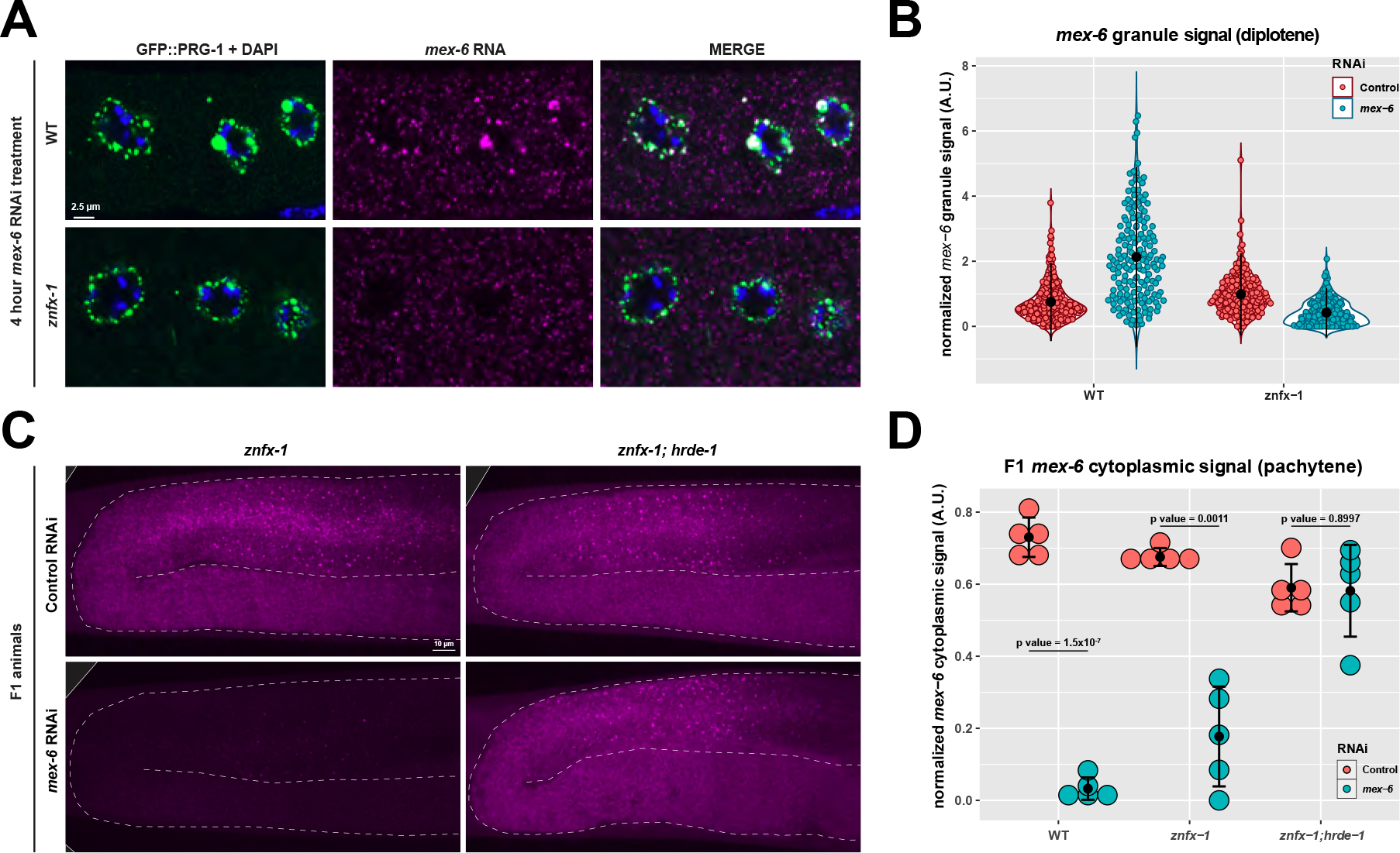
*znfx-1* is required for enrichment of RNAi-targeted transcripts in germ granules. A) Photomicrographs of oocytes in wild-type and *znfx-1* mutant animals after 4 hours of RNAi treatment showing the germ granule marker GFP::PRG-1 (green), DNA stained with DAPI (blue), and *mex-6* RNA (magenta). B) Graph comparing the mean *mex-6* RNA FISH signal in germ granules in wild-type and *znfx-1* mutant animals after 4 hours of RNAi treatment. Each dot represents an individual granule. Central black dot and error bars represent the mean and standard deviation, respectively. Five worms were quantified for each condition. Values (arbitrary units) were normalized to *puf-5* RNA FISH signals visualized in the same granules (Methods). C) Maximum projection photomicrographs of germlines showing *mex-6* RNA (magenta) in F1 progeny of animals with the indicated RNAi treatment. D) Graph comparing the mean *mex-6* RNA FISH signal in germ granules in the pachytene region of *znfx-1* and *znfx-1; hrde-1* F1 progeny derived from animals with the indicated RNAi treatment. Each dot represents a single worm. Central black dot and error bars represent the mean and standard deviation, respectively. Values (arbitrary units) were normalized to *puf-5* RNA FISH signals visualized in the germlines (Methods). P values were calculated using an unpaired t-test.

We detected an increase in the intensity distribution of *mex-6* nuclear puncta in *znfx-1* mutants at the 4-hour time point, earlier than in wild-type (8 hour) (Fig S5A,SB). This premature peak in nuclear signals was followed by a subsequent decrease to levels lower than the non-RNAi condition by the 24-hour time point in *znfx-1* mutants (Fig S5A,B). No changes in nuclear signal were observed in *znfx-1; hrde-1* double mutant animals, indicating that the nuclear response in *znfx-1* mutants was dependent on *hrde-1*, as in wild-type (Fig S5C,S5D). We conclude that *znfx-1* is required for robust recruitment of *mex-6* transcripts to nuage in P0 animals but is not required for RNA degradation in the cytoplasm or for engagement of the nuclear RNAi machinery in P0 animals. Despite a failure to silence the *mex-6* locus and to enrich *mex-6* transcripts in nuage, *znfx-1; hrde-1* P0s still showed rapid loss of cytoplasmic *mex-6* RNA throughout the germline, confirming that neither ZNFX-1 nor HRDE-1 is required for RNA turn over in the cytoplasm of P0 animals (Fig S5C).

In *znfx-1* F1 animals, we observed a partial (∼50%) reduction in cytoplasmic accumulation of *mex-6* transcripts in the pachytene rachis and no accumulation in nuage in the pachytene region (Fig 5C,5D,S5E). The intensity distribution of nuclear puncta was reduced as observed in wild-type F1s (Fig S5B). This reduction was dependent on *hrde-1,* as nuclear puncta intensities of *znfx-1; hrde-1* F1s matched that of the no-RNAi controls (Fig S5C, S5D). We conclude that *znfx-1* is not required for silencing of the locus in P0 and F1 animals (nuclear response) but is required for the accumulation of targeted transcripts in nuage in P0 and F1 animals (cytoplasmic response).

### *hrde-1* and *znfx-1* are required additively for maximal silencing in F1 animals

Unlike in P0 animals, cytoplasmic *mex-6* RNA levels in *znfx-1; hrde-1* F1s were indistinguishable from no-RNAi controls, indicating that *znfx-1* and *hrde-1* are both required for maximal silencing in F1 animals (Fig 5C, 5D). To examine this further, we compared *mex-6* RNA levels using RT-PCR in wild-type, *znfx-1, hrde-1,* and *znfx-1; hrde-1* double mutants F1 animals (Fig S6A). These experiments confirmed partial silencing of *mex-6* transcripts in the single mutants, and complete loss of silencing in the double mutant (Fig S6A). We obtained similar results when targeting two other germline-expressed genes by RNAi (*oma-1* and *puf-5*) (Fig S6B, S6B). We conclude that *hrde-1* and *znfx-1* contribute independently to silencing in F1s and are required additively for maximal silencing.

### *hrde-1* and *znfx-1* are responsible for distinct populations of sRNAs in F1 progeny

The additive phenotype of the *znfx-1; hrde-1* double mutant suggested that *hrde-1* and *znfx-1* function in separate mechanisms to maintain nuclear and cytoplasmic silenced states. To examine this possibility further, we sequenced sRNAs in wild-type, *hrde-1*, *znfx-1* and *znfx-1; hrde-1* F1 adult progeny of *mex-6* fed P0s. As expected, wild-type F1s exhibited a 23-fold increase in sRNAs mapping to the *mex-6* locus compared to no-RNAi controls, with a dominant peak corresponding to the location targeted by the dsRNA trigger fed to the P0 generation (Fig 6A, 6B). A similar pattern was observed in *hrde-1* mutants, although the overall increase was roughly only 83% that observed in wild-type (Fig 6A, 6B). In contrast, *znfx-1* mutants only showed a modest increase in sRNAs corresponding to 6% that of wild-type (Fig 6A, 6B).

**Figure 6.**
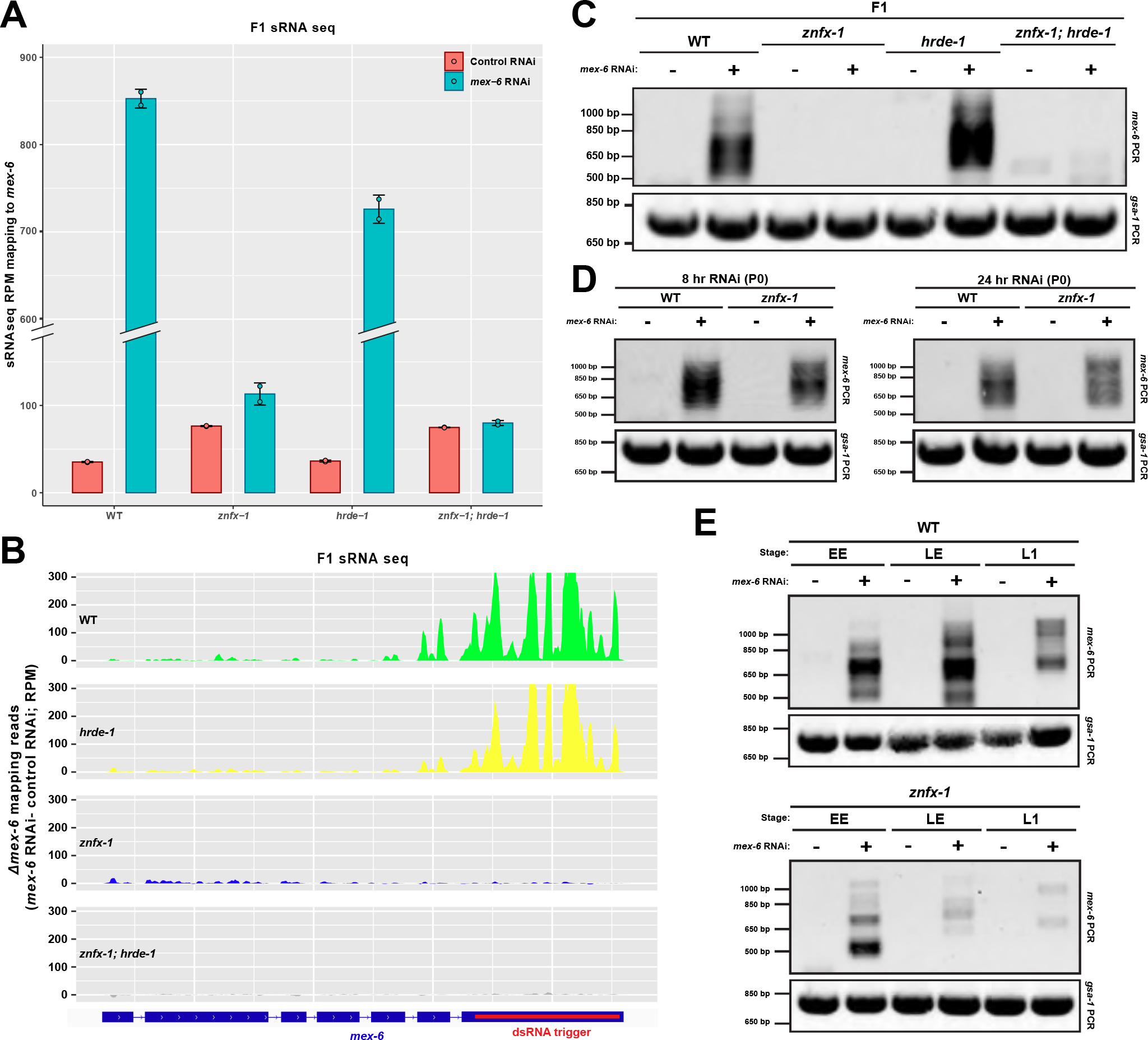
ZNFX-1 and HRDE-1 function in separate pathways contributing to RNAi inhertiance. A) Bar graph depicting the number of sRNAseq reads (normalized per million) mapping to the *mex-6* transcript in wild-type, *znfx-1*, *hrde-1*, and *znfx-1; hrde-1* mutant F1 progeny derived from animals with the indicated RNAi treatment. Each dot represents a technical replicate and error bars represent standard deviation between the respective replicates. B) Genome browser view of the sRNAseq reads mapping to the *mex-6* locus in wild-type, *hrde-1*, *znfx-1*, and *znfx-1; hrde-1* mutant F1 progeny of animals treated with *mex-6* RNAi. sRNA reads were binned across the *mex-6* gene, and the number of in each reads in each bin under the control RNAi condition were subtracted from the number of reads in each respective bin under the *mex*-6 condition. Positioning of the dsRNA trigger administered in the P0 generation is indicated in red. C)Gel showing PCR amplification of pUGylated *mex-6* RNA from lysates derived from F1 progeny from animals of the indicated genotype treated with control (“-“) or *mex-6* (“+”) RNAi (top panel). The *gsa-1* transcript has a genomically-encoded 18-nue poly(UG) stretch and is used here as a positive control for pUG amplification (bottom panel; Shukla et al., 2020). D)Gel showing PCR amplification of pUGylated *mex-6* RNA from lysates derived from P0 animals of the indicated genotype and treated for 8 hours with control (“-“) or *mex-6* (“+”) RNAi (top panel). *gsa-1* is the pUG amplification control (bottom panels). E) Gel showing PCR amplification of pUGylated *mex-6* RNA from lysates derived from early embryos (EE), late embryos (LE), and first larval stage F1 progeny from animals of the indicated genotype treated with control (“-“) or *mex-6* (“+”) RNAi. *gsa-1* is the pUG amplification control (bottom panels).

Strikingly, the distribution of sRNAs in *znfx-1* mutants showed no preference for the region targeted by the trigger (Fig 6B). Instead sRNAs appeared distributed throughout the *mex-6* locus, with a slight bias for the 5’ end of the transcript. Consistent with the complete lack of inherited RNAi response, *znfx-1; hrde-1* double mutants exhibited no significant differences in sRNAs across the *mex-6* locus in *mex-6* RNAi vs. control RNAi conditions (Fig 6A, 6B). These observations suggest that *hrde-1* and *znfx-1* are required for the amplification of distinct pools of sRNAs across the *mex-6* locus, with *znfx-1* being required for the majority of sRNA generation, especially around the sequence targeted by the original trigger. We noticed that the number of sRNA reads mapping to the *mex-6* locus in *znfx-1* and *hrde-1* single mutants added up to only 89% of the reads observed in wild-type F1s (Fig S6G; see Methods). This observation confirms that the ZNFX-1 and HRDE-1 amplification cycles function mostly independently, with possibly some synergy between the two cycles accounting for ∼10% of sRNAs observed in wild-type.

To determine whether *znfx-1* is also required for sRNA amplification in P0 animals, we sequenced sRNAs in wild-type and *znfx-1* hermaphrodites at different time points following feeding onset. We observed ∼200-fold increase in sRNA accumulation at the *mex-6* locus in wild-type and *znfx-1* P0 animals compared to no-RNAi conditions (Fig S6D). The increase in sRNAs levels in *znfx-1* mutants were slightly lower than in wild-type (∼16% reduction), suggesting that *znfx-1*, although not essential, contributes modestly to sRNA amplification in P0s. In contrast, in F1 progeny*, znfx-1* is required for the majority of sRNA production, especially in the region corresponding to the trigger (Fig. 6A-B).

### *znfx-1,* but not *hrde-1*, is required for sustained accumulation of pUGylated transcripts in F1 progeny

sRNAs mapping to the trigger region can be derived directly by Dicer cleavage of the dsRNA trigger (1° sRNAs) or from RdRPs that use pUGylated transcripts as templates to generate 2° sRNAs (Shukla et al., 2020, 2021). To determine whether *hrde-1* or *znfx-1* are required for pUGylation of the *mex-6* RNA, we amplified pUGylated *mex-6* transcripts from RNA extracted from wild-type, *hrde-1*, *znfx-1*, and *znfx-1; hrde-1* F1 animals. As expected, wild-type F1 animals exhibited abundant pUGylated *mex-6* transcripts (Fig 6C). pUGylated *mex-6* transcripts were also observed in *hrde-1* F1 animals (Fig 6C). In contrast no pUGylated *mex-6* transcripts were observed in *znfx-1* or *znfx-1; hrde-1* F1 adult animals (Fig 6C), indicating that ZNFX-1 is required for accumulation of pUGylated mRNAs in F1 adult progeny. Similar results were observed in the F1 progeny of *puf-5* and *oma-1* RNAi-ed P0s (Fig S6E, S6F).

To determine whether *znfx-1* is also required for pUGylation in animals exposed to the dsRNA trigger, we repeated the pUGylation assays on *znfx-1* P0 worms at 8 and 24 hours after RNAi exposure. We observed robust accumulation of pUGylated *mex-6* transcripts, indicating that *znfx-1* is not required for the initial production of pUGylated RNAs in animals exposed to the dsRNA trigger (Fig 6D). We also detected pUGylated mRNAs in *znfx-1* F1 embryos (Fig 6E). We conclude that *znfx-1* is not required for pUGylation in P0 animals or for transfer of pUGylated transcripts to F1 embryos. ZNFX-1, however, is required for sustained production of pUGylated RNAs in adult F1 animals.

### ZNFX-1 associates with pUGylated *mex-6* transcripts and is required for accumulation of pUGylated RNAs in germ granules

ZNFX-1 immunoprecipitates with transcripts targeted by RNAi (Wan et al., 2018). To determine whether ZNFX-1 interacts with pUGylated transcripts, we immunoprecipitated FLAG-tagged ZNFX-1 in animals exposed to *puf-5* or *mex-6* RNAi triggers for 12 hours. We performed reverse transcription reactions on the immunoprecipitates using a poly(UG) specific RT primer to amplify pUGylated RNAs (Shukla et al., 2020, 2021). We found that ZNFX-1 co-immunoprecipitated with *mex-6* pUGylated transcripts in animals exposed to the *mex-6* trigger and with *puf-5* pUGylated transcripts in animals exposed to the *puf-5* trigger (Fig 7A, S7A, S7B). We conclude that ZNFX-1 is in a complex with pUGylated RNAs.

**Figure 7:**
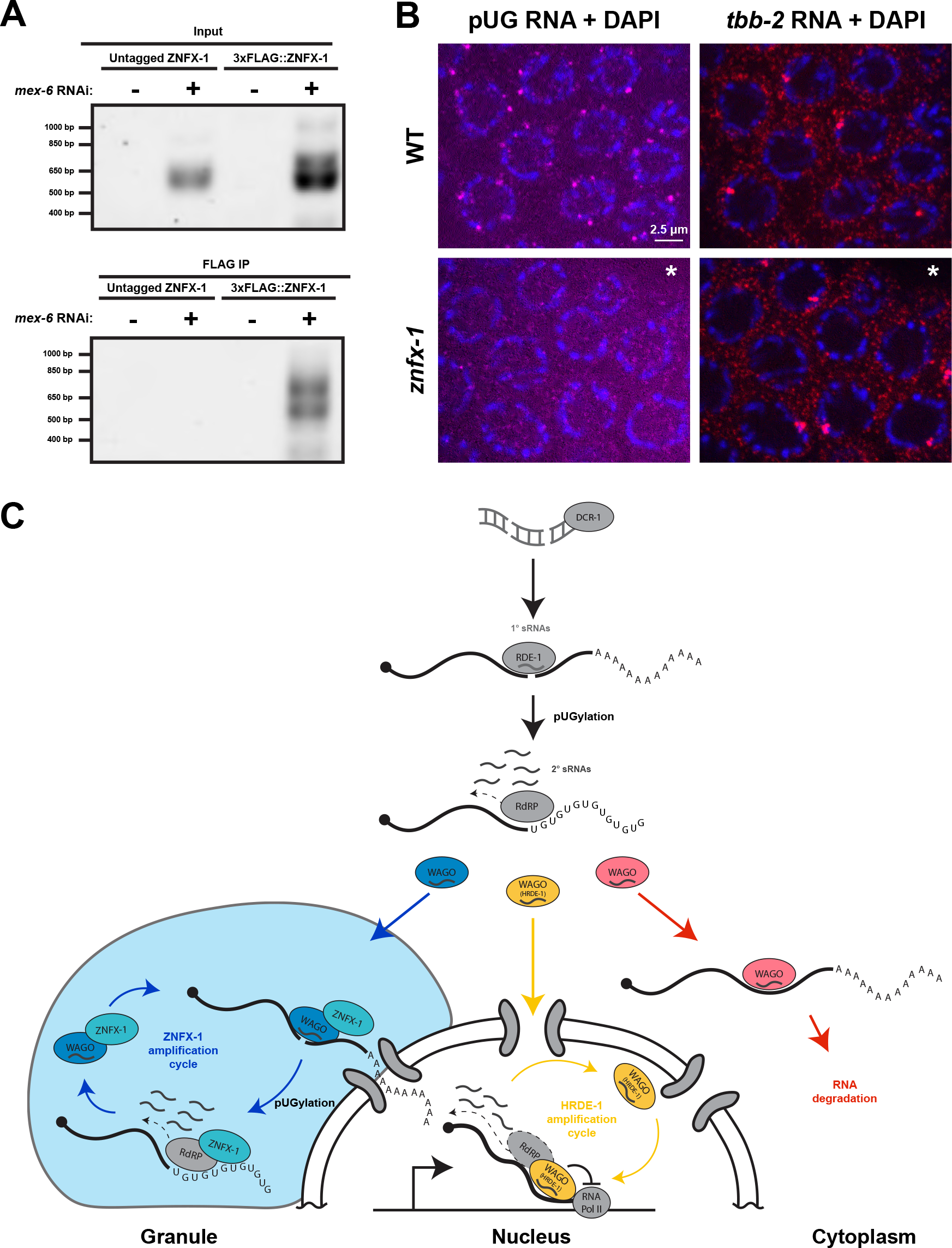
ZNFX-1 immunoprecipitates with pUGylated RNAs and is required for localization of pUGylated RNAs to germ granules. A) Gel showing PCR amplification of pUGylated *mex-6* RNA from input (top panel) or FLAG immunoprecipitates (bottom panel) from animals where the *znfx-1* locus or tagged with 3xFLAG. Lysates were collected from adult worms grown for 12 hours on either *mex-6* (“+”) or *puf-5* (“-”) RNAi. B) Photomicrographs of pachytene nuclei in dissected germline from wild-type or *znfx-1* mutant animals showing endogenous pUGylated RNAs (magenta), control red), and DNA stained with DAPI (blue). Images marked with asterisks were enhanced for contrast for visualization purposes. See Fig. S7C for evenly mages. C)Model for exogenous RNAi. dsRNA is processed into primary sRNAs that load with RDE-1 to target complementary transcripts for pUGylation. pUGylated transcripts recruit RdRPs to generate secondary sRNAs in the cytoplasm. Secondary sRNAs bind to secondary Argonautes (“WAGOs”), which initiate three distinct first pathway leads to degradation of cytoplasmic transcripts with no further sRNA amplification. On its own, this pathway is sufficient to silence gene in animals exposed to dsRNA triggers but is not sufficient to propagate the RNAi response across generations. A second pathway dependent on the onaute HRDE-1 partially silences the locus and uses nascent transcript as templates for the production of “tertiary” sRNAs. A third pathway dependent e helicase ZNFX-1 enriches targeted transcripts in nuage where they are pUGylated and used as templates for further “tertiary” sRNAs amplification. As feedback into their respective cycles ensuring inheritance of the silenced state. Possible cross talk between the HRDE-1 and ZNFX-1 cycles is not Discussion for further considerations.

Animals not exposed to exogenous RNAi triggers naturally contain pUGylated transcripts, due to targeting by endogenous sRNAs (Shukla et al., 2020). Endogenous pUGylated transcripts can be visualized by FISH using a poly(AC) probe and have been reported to accumulate in nuage (Shukla et al., 2020). Consistent with this report, we detected endogenous pUGylated transcripts in nuage in the pachytene region and in oocytes (Fig 7B, S7C, S7D). Remarkably, in *znfx-1* mutant germlines, accumulation of pUGylated transcripts in nuage was strongly reduced in both the pachytene region and in oocytes (Fig 7B, S7C, S7D). We conclude that ZNFX-1 is required for accumulation (and/or possibly synthesis) of endogenous pUGylated RNAs in nuage.

## Discussion

In this study, we have used fluorescent *in situ* hybridization, small RNA sequencing and pUGylation assays to examine the fate of mRNAs targeted for silencing by feeding RNAi. Together with prior studies (referenced below), our findings suggest the following model for silencing by an exogenous dsRNA trigger (Fig. 7C). Primary siRNAs derived from the double-stranded RNA trigger load onto the primary Argonaute RDE-1. RDE-1 recognize complementary transcripts and marks them for cleavage, pUGylation, and synthesis of secondary sRNAs by RNA-dependent RNA polymerases (Tabara et al., 1999; Sijen et al., 2001; Yigit et al., 2006; Sijen et al., 2007; Pak et al., 2007; Tsai et al., 2015; Shukla et al., 2020). Secondary sRNAs are loaded onto secondary Argonautes (HRDE-1 and other WAGOs (Yigit et al., 2006; Gu et al., 2009; Ashe et al., 2012; Buckley et al., 2012; Shirayama et al., 2012)) that, in turn, activate three parallel silencing pathways. In the first pathway, WAGOs target transcripts in the cytoplasm for rapid degradation by an unknown mechanism. In the second pathway, HRDE-1 shuttles into the nucleus to initiate “nuclear RNAi”, a silencing program that suppresses but does not eliminate transcription of the locus. In the third pathway, WAGOs that associate with ZNFX-1 recruit a subset of targeted transcripts to germ granules and initiate a new cycle of pUGylation and sRNA amplification. Only the HRDE-1 and ZNFX-1 cycles generate “tertiary” sRNAs that feedback into their respective cycles to generate parallel, self-reinforcing sRNA amplification loops. The HRDE-1 and ZNFX-1 amplification loops are transmitted to the next generation independently of each other and both are required for maximum silencing in F1 progeny. In the following sections, we summarize evidence supporting the three silencing pathways and discuss remaining open questions.

### Pathway 1: RDE-1 and MUT-16 dependent amplification of secondary sRNAs induces RNA degradation in the cytoplasm

Under our feeding RNAi conditions, we detected a reduction in transcript levels in the cytoplasm after 4 hours of feeding, eventually reaching 95% reduction by 24 hours. Loss of transcripts was most rapid in growing oocytes, consistent with prior reports showing that trigger RNAs enter the germline in a manner dependent on the oocyte yolk receptor RME-2 (Marré et al., 2016; Wang & Hunter, 2017). As expected (Tabara et al., 1999; Zhang et al., 2011; Phillips et al., 2012), mRNA degradation was dependent on the primary Argonaute RDE-1 and on MUT-16, a scaffolding protein required for amplification of secondary sRNAs.

mRNAs targeted by miRNAs for degradation have been reported to enrich in P bodies (Liu et al., 2005; Shih et al., 2011), RNA granules in the cytoplasm that contain components of the RNA degradation machinery (Luo et al., 2018). In our feeding experiments, we observed enrichment of targeted mRNAs in clusters that overlapped with nuage components, but this enrichment did not appear linked to degradation. Most strikingly, no nuage enrichment was observed in *znfx-1* mutants despite normal RNA degradation in these animals. We conclude that, unlike miRNA-induced RNA degradation, RNAi-induced RNA degradation does not require visible enrichment of RNA in cytoplasmic granules. The robust sRNA amplification observed in *znfx-1* P0 animals also suggests that secondary sRNA amplification initiated by primary sRNAs occurs in bulk cytoplasm or at a minimum does not require accumulation of targeted transcripts in granules. We cannot exclude, however, that transit through nuage or some other RNA granules, in the absence of visible accumulation, is required for secondary sRNA amplification and/or RNA degradation. The RDE-1-initiated cycle of pUGylation and sRNA amplification is sufficient to eliminate most cytoplasmic transcripts in animals exposed to the dsRNA trigger. This cycle is not self-perpetuating, however, and on its own will eventually self-extinguish leaving no memory of the RNAi response.

### Pathway II: HRDE-1-dependent silencing targets nascent transcripts within hours of dsRNA exposure and reduces, but does not eliminate, transcription of the targeted locus

Quantification of FISH signals at the targeted locus revealed a transient increase in nascent transcripts starting at 6 hours of feeding, followed by a subsequent decrease in signal at 24 hours in P0 animals, and even lower levels in F1 animals. This response requires the Argonaute HRDE-1, a component of the nuclear RNAi pathway. We speculate that the initial increase in FISH signal reflects stalling of RNA polymerase II and/or stalling of pre-mRNA processing causing nascent transcripts to accumulate at the locus. Stalling of RNA polymerase has previously been implicated in nuclear RNAi (Guang et al., 2010; Liao et al., 2020).

Additionally, several lines of evidence have suggested a close connection between RNAi and splicing, including apparent co-evolution of the RNAi and splicing machineries (Tabach et al., 2013), splicing factors identified as HRDE-1 interactors (Akay et al., 2017; Tyc et al., 2017), sRNA defects associated with mutations or knock down of spliceosome components (Kim et al., 2005; Newman et al., 2018), and insensitivity to nuclear RNAi of an endogenous transcript whose introns were removed by genome editing (Wan et al., 2020). It will be interesting to determine which components of the nuclear RNAi machinery in addition to HRDE-1 are required for the transient increase in nascent transcripts at the locus and whether a slow-down in RNA polymerase elongation or splicing is primarily responsible.

A recent study (Yang et al., 2021) also reported a transient increase in RNA signal in the pachytene region early in the RNAi response but interpreted this observation differently. In their analysis, using *oma-1* as a gene model, they reported that the bright RNA foci overlapped with a single enlarged perinuclear nuage condensate (P granule) outside each pachytene nucleus. Although we did observe targeted transcripts in nuage (see below), they did not concentrate in just one condensate, nor did we observe enlargement of a single P granule per nucleus under live or fixed conditions (Fig. S8A-B). In our hands, the bright foci did not correspond to P granules but rather overlapped with DAPI at the nuclear periphery and often could be resolved in two or more closely apposed dots (<150nm; Figs. S1, S8C), consistent with nascent transcripts at the locus, which at this stage of germline development is present in four tightly synapsed copies. In their analyses, Yang et al. reported that the *oma-1* locus and associated nascent transcripts appear as two well-separated (∼0.5um) foci in the center of pachytene nuclei that eventually move to the periphery and merge near the enlarged P granule during the RNAi response. These observations were based on a new DNA/RNA hybridization protocol and contrast with several studies using standard DNA *in situ* methods that have demonstrated that, throughout the pachytene region, homologous chromosomes are tightly synapsed at the nuclear periphery and excluded from a central, DNA-depleted zone occupied by the nucleolus (Phillips and Dernburg, 2006; Dernburg et al., 1998; Lui & Colaiácovo, 2013; Macqueen et al., 2002; Zalevsky et al., 1999).

At 24 hours post-feeding, we observed a decrease in the accumulation of nascent transcripts in the pachytene region, which became even more acute in F1 animals. We speculate that the decrease reflects a reduction in transcription initiation at the locus. The nuclear RNAi machinery deposits chromatin marks at the locus predicted to decrease transcription (Burkhart et al., 2011; Burton et al., 2011; Gu et al., 2012; Mao et al., 2015; Schwartz-Orbach et al., 2020). We note, however, that despite the apparent decrease in transcriptional output, we continued to observe transcripts in perinuclear nuage even in F1 animals, indicating that a baseline level of transcription and export is maintained at the silenced locus. In *S. pombe,* transcription is also maintained at the silent locus, but export is blocked and replaced by rapid degradation of nuclear transcripts (Martienssen & Moazed, 2015). We speculate that this difference reflects a *C. elegans*-specific adaptation that allows mature transcripts to be used as templates for sRNA amplification in perinuclear condensates (see Pathway III).

sRNA sequencing analyses suggest that the HRDE-1 cycle generates sRNAs that map throughout the locus without preference for the trigger area. Interestingly, HRDE-1-dependent sRNAs exhibit a slight preference for the 5’ end of the transcript. A similar pattern was described previously in the context of transgenes and endogenous transcripts targeted by endogenous sRNA pathways and also was found to be dependent on the nuclear RNAi machinery (Sapetschnig et al., 2015). One hypothesis is that the 5’ bias is due to RNA-dependent RNA polymerases that use nascent transcripts as templates for sRNA synthesis as described in *S. pombe*. Consistent with this hypothesis, the nuclear RNAi machinery has been shown to interact with pre-mRNAs at the locus, which naturally exhibit a 5’ bias (Burkhart et al., 2011; Guang et al., 2010). We suggest that HRDE-1, initially loaded with secondary sRNAs templated in the cytoplasm, initiates a nuclear cycle of sRNA amplification by recruiting an RdRP to nascent transcripts. The HRDE-1 cycle generates “tertiary” sRNAs which in turn become complexed with HRDE-1 to perpetuate the cycle. The RNA-dependent RNA polymerase (RdRP) EGO-1 has been reported to localize within nuclei (Claycomb et al., 2009), but a specific molecular interaction between EGO-1 and HRDE-1 has not been reported. Analyses of silencing in operons, however, has provided indirect evidence for an RdRP activity in nuclei. Operons are loci that generate long primary transcripts that are cleaved in the nucleus into multiple transcripts by trans-splicing (Agabian, 1990). Several studies have documented that sRNA production initiated on a specific transcript spreads to other transcripts in the same operon, implying that nascent transcripts are used as templates for sRNA production before trans-splicing (Bosher et al., 1999; S. Guang et al., 2008; Ouyang et al., 2019; Sapetschnig et al., 2015). Although we favor a model where HRDE-1 and associated machinery use nascent transcripts to direct sRNA synthesis (Fig. 7C), we cannot exclude the possibility that HRDE-1-dependent sRNA amplification actually occurs outside of the nucleus. For example, HRDE-1 could deposit marks on nascent transcripts that target them for sRNA amplification after export into the cytoplasm. Investigation into the factors that support HRDE-1-dependent sRNA production is an important future goal.

### Pathway III: ZNFX-1 memorializes the dsRNA trigger by initiating a self-perpetuating pUGylation/sRNA amplification cycle, likely within nuage

Our genetic analyses indicate that the HRDE-1 amplification cycle is not sufficient for maximum silencing in F1 progeny. A second cycle dependent on the nuage protein ZNFX-1 is also required. Unlike the HRDE-1 cycle, the ZNFX-1 cycle generates sRNAs focused primarily on the area of the transcript targeted by the original trigger. Consistent with that observation, ZNFX-1 (but not HRDE-1) is required to maintain the production of pUGylated transcripts in adult F1s. Interestingly, ZNFX-1 is NOT required for the initial production of pUGylated transcripts in P0 animals exposed to the trigger. These observations suggest that ZNFX-1 becomes essential for pUGylation when the dsRNA trigger and primary sRNAs become limiting. One possibility is that ZNFX-1 extends the half-life of pUGylated mRNAs by protecting them from being targeted for degradation by secondary Argonautes. Additionally, ZNFX-1 could also function as a bridge between secondary Argonautes and the pUGylation machinery, allowing secondary Argonautes to generate new pUGylated transcripts for synthesis of “tertiary” sRNAs. Consistent with this model, ZNFX-1 has been reported to immunoprecipitate in complexes that also contain the secondary Argonautes WAGO-1 and WAGO-4 and the RNA-dependent RNA polymerase EGO-1 (Barucci et al., 2020; Ishidate et al., 2018; Wan et al., 2018) and we show here that ZNFX-1 complexes also contain pUGylated transcripts. We suggest that tertiary sRNAs generated in the ZNFX-1 loop feedback into additional cycles of pUGylation and sRNA amplification to ensure propagation of sRNA amplification across generations. Because this self-perpetuating cycle is initiated by secondary sRNAs that target the trigger region, the ZNFX-1 cycle “memorializes” the position of the trigger. A role for ZNFX-1 in a self-propagating sRNA amplification cycle is consistent with the role proposed for Hrr1, the *S. pombe* ortholog of ZNFX-1, which functions in a nuclear complex with an RNA-dependent RNA polymerase to amplify sRNAs off nascent transcripts (Motamedi et al., 2004). Unlike Hrr1 which has been shown to be nuclear (Motamedi et al., 2004), ZNFX-1 is prominent in nuage (Ishidate et al., 2018; Wan et al., 2018) and is required to concentrate targeted transcripts (and pUGylated RNAs) to nuage (this work). We speculate therefore that *C. elegans* ZNFX-1 functions outside of the nucleus on mature transcripts exported from the nucleus into nuage.

It has been suggested that, in *C. elegans*, initiation of sRNA amplification by non-primary sRNA/Argonaute complexes is limited *in vivo* to prevent dangerous run-away loops (Pak et al., 2012). We speculate that enrichment of ZNFX-1 in nuage serves to place the ZNFX-1 amplification loop under tight regulation by competing sRNA pathways (such as the piRNA pathway) that protect transcripts from permanent silencing (Shukla et al., 2021). Consistent with this, loss of the nuage component PRG-1 causes indefinite silencing and pUGylation of RNAi-silenced transcripts (Shukla et al., 2021). Enrichment of pUGylated transcripts in nuage may also serve to protect them from degradation in the cytoplasm. By maintaining a transcript pool in nuage, ZNFX-1 prevents self-extinction of the RNAi response that might arise as a consequence of rapid transcript turn over in the cytoplasm. ZNFX-1 homologs in mice and humans function in the primary immune response against RNA viruses and bacteria (Wang et al., 2019; Le Voyer et al., 2021; Vavassori et al., 2021). We do not yet know how these functions relate to those described here for *C. elegans* ZNFX-1 and described previously for *S. pombe* Hrr1 (Motamedi et al., 2004). We speculate that a common function for ZNFX-1 homologs across eukarya may be to memorialize transcripts for long-term silencing.

### The HRDE-1 and ZNFX-1-dependent amplification loops contribute mostly additively to RNAi inheritance

Our findings suggest that inheritance of the silenced state by F1 progeny not exposed to the trigger involves two pathways: 1) the HRDE-1 “nuclear” sRNA amplification pathway that generates sRNAs across the locus and does not appear to involve pUGylation, and 2) the ZNFX-1 “nuage” pathway that uses cycles of pUGylation and sRNA amplification to memorialize the dsRNA trigger. In contrast to RDE-1-initiated sRNA amplification and RNA degradation in P0 animals exposed to the trigger, the HRDE-1 and ZNFX-1 programs are self-sustaining cycles that maintain a pool of targeted transcripts for use as templates for sRNA amplification. Our analyses suggest that both the HRDE-1 and ZNFX-1 programs are required for full silencing in F1 animals. Because our analyses were restricted to feeding RNAi against three loci with similar expression patterns in the germline (*mex-6*, *puf-5* and *oma-1*), it is possible that reliance on the HRDE-1 or ZNFX-1 programs will vary between different loci and in response to different silencing triggers, such as endogenous sRNAs. Consistent with this hypothesis, different genetic requirements for RNAi inheritance have been documented for *oma-1* and a germline expressed *gfp::H2B* transgene (Kalinava et al., 2017; Lev et al., 2019).

Although our genetic analyses suggest that the HRDE-1 and ZNFX-1 pathways function primarily independently of each other, two lines of evidence hint at possible cross-talk. First, we found that the sum of *mex-6* sRNAs induced by RNAi in *hrde-1* and *znfx-1* F1s adds up to only 89% of what is observed in wild-type. While this observation will need to be repeated in different contexts to ensure reproducibility, it raises the possibility that sRNAs produced by one amplification cycle extend sRNA production in the other cycle. Second, we observed an accelerated nuclear RNAi response in *znfx-1* P0 animals compared to wild-type, suggestive of competition between the ZNFX-1 and HRDE-1 pathways. One possibility is that in the absence of ZNFX-1, more secondary sRNAs and/or RdRPs are available to fuel the nuclear pathway in the early stages of the RNAi response. Alternatively, ZNFX-1 may antagonize HRDE-1-initiated transcriptional silencing to ensure sufficient production of mature mRNAs for use in the ZNFX-1 cycle. More complex interplays involving Argonautes that participate in multiple sRNA amplification mechanisms are also possible. How the RDE-1, HRDE-1, and ZNFX-1-driven sRNA amplification mechanisms (Fig. 7C) are coordinated in cells and across generations will be an important focus for future investigations.

## Author contributions

Experiments were conducted by JPTO and WZ. Experimental design and analysis were conducted by JPTO, WZ and GS. JPTO and GS wrote the manuscript.

## Acknowledgments

We thank Scott Kennedy, John Kim, Antony Jose, and Craig Mello for sharing expertise and unpublished data, the Johns Hopkins Integrated Imaging center (S10OD023548) for microscopy support, and the Johns Hopkins University School of Medicine Genetic Resources Core Facility for sequencing support. We also thank the Seydoux lab, Baltimore Worm Club, Tatjana Trcek, Jeff Corden, Taylor Swift and Justinian for their support during this project. Funding was provided by the National Institutes of Health (GS: Grant number R37HD037047; JPTO: T32GM007445 and F31HD103428) and a JHU Provost Award to WZ. GS is an investigator of the Howard Hughes Medical Institute. 

## Methods

### Strains and maintenance

All strains were cultured/maintained at 20° C on either OP50 bacteria plated on NNGM media or NA22 bacteria plated on Enriched Peptone (EP) media. The following strains were used in this study: N2 (JH1), *znfx-1(gg561) II* (YY996; Wan et al., 2018), *hrde-1(tm1200) III* (YY538; Buckely et al., 2012), *znfx-1(gg561) II; hrde-1(tm1200) III* (JH4054; this study), *prg-1(ne4523[gfp::tev::flag::prg-1]) I* (WM527; Shen et al., 2018), *znfx-1(gg544[3xflag::gfp::znfx-1])* (YY916; Wan et al., 2018), *prg-1(ne4523) I; znfx-1(gg561) II* (JH4055; this study), *prg-1(ne4523) I; hrde-1(tm1200) III* (JH4056; this study), *prg-1(ne4523) I; znfx-1(gg561) II; hrde-1(tm1200) III* (JH4057; this study), *prg-1(ne4523) I; rde-1(ne219) V* (JH4058; this study)*, prg-1(ne4523) mut-16(pk710) I* (JH4059; this study), *ego-1(ne4518[gfp::ego-1]) I; znfx-1(ne4355[3Xflag::tev::znfx-1])II* (WM514; Ishidate et al., 2018), *ego-1(ne4518[gfp::ego-1]) I* (WM522; Ishidate et al., 2018), *glh-1(ax3843[glh-1::eGFP]) I* (JH4022; Paix et al., 2016).

### RNA extraction and purification

RNA extraction was performed on up to 100 uL of worms flash frozen in liquid nitrogen and stored at -80° C. First, frozen samples were resuspended in 1 mL of Trizol (ThermoFisher; Cat #: 15596026) and subjected to three freeze-thaw cycles. Samples were then shaken at 1500 RPM for five minutes at RT in a benchtop shaker (Benchmark Scientific; Model #: H5000-HC). Samples were then incubated for 5 minutes at RT with no shaking. 200 uL of chloroform was then added to each tube, and the samples were then shaken by hand for 15 seconds, followed by a 2-3 minute incubation at RT. Tubes were then spun at 4° C at 12,000xg for 15 minutes. The upper aqueous phase was then removed and an equal volume of 95-100% ethanol was added and mixed. These samples were then input into the Zymo RNA Clean & Concentrator Kit columns (Zymo; Cat #: R1017) and purified according to the kit manual. On-column DNase I digestions with MgCl2 buffer (ThermoFisher; Cat # EN0521) were used for each RNA prep to remove contaminating DNA. Samples were eluted in water.

### Plasmid construction

RNAi plasmids were constructed using the In-Fusion® HD Cloning Kit (Takara Bio; Cat #: 639650) and the L4440 vector. Primers for cloning the 3’ portion of *mex-6* (600 nt), *puf-5* (500 nt), and *oma-1* (600 nt) were designed using the Takara Bio In-Fusion Cloning online design tool and are listed in Table S1. NEB Phusion PCRs (NEB; Cat #: M0531S) were conducted from reverse transcriptase reactions generated from the SuperScript™ VILO™ cDNA Synthesis Kit (Thermofisher; Cat #: 11754050) and RNA extracted from adult animals (see RNA extraction). The L4440 plasmid was digested using XbaI and NcoI, gel purified, and added to the In-Fusion® HD Cloning reaction along with the amplified PCR products. Reactions were then transformed into Stellar™ competent cells (Takara Bio; Cat #: 636766), and plasmids were subsequently isolated using Qiagen mini-prep kits (Qiagen; Cat #: 27104).

### RNAi assays

RNAi plates were made as follows: RNAi constructs were transformed into HT115 bacteria, and overnight starter cultures were then grown from the HT1115 transformants in 100 ug/mL ampicillin LB liquid media culture at 37° C with vigorous shaking. The starter culture was then used to inoculate a fresh LB/ampicillin culture at a ratio of 1:100 (e.g. 10 mLs starter culture into 990 mLs LB), and grown for 6 hours with vigorous shaking at 37° C. RNAi cultures were then induced with IPTG for a final concentration of 500 uM and shaken for 30 more minutes at 37° C. Cultures were then spun down and resuspended in 1/20 of the culture volume with 100 ug/mL ampicillin/500 uM IPTG LB (e.g. 50 mLs for 1000 mLs of culture) and densely plated onto NNGM agar containing 100 ug/mL carbenicillin and 1 mM IPTG. Plates were then allowed to dry, and then subsequently used for RNAi.

RNAi assays were conducted as follows: embryos were isolated from gravid mothers through bleaching and placed in M9 overnight at 20° C and 110 RPM shaking. Synchronized L1s were then plated onto NA22 bacteria grown on EP media. Worms were then collected at the adult stage approximately 60 hours after plating and washed and collected using a filter. Worms were then plated onto either control or gene-specific RNAi plates for the specified lengths of time, and then washed and collected for fixation (see FISH assays) or RNA collection (see RNA collection).

For RNAi inheritance assays, synchronized P0 L1s were plated onto NA22 plates for approximately 60 hours and were collected as adults with a filter. Worms were then put onto either control or *mex-6* RNAi plates for 24 hours. After 24 hours, adult worms were collected, and embryos were isolated through bleaching. Embryos were further synchronized by shaking overnight (110 RPM) at 20° C. The worms were then plated onto NA22 plates and collected for fixation/RNA extraction ∼72 hours later. For examining the F2 generation, F1 adult worms were bleached, and embryos were isolated. F2 adults were collected ∼72 hours after plating synchronized L1s onto NA22 plates.

For experiments examining late vs early F1 embryos, adult worms fed RNAi for 24 hours were isolated. Embryos extracted from bleaching mothers were considered “early embryos.” Embryos collected from the plate were considered “late embryos.” Late embryo samples were bleached to avoid any hatched L1s also being collected.

### FISH protocol

For whole worm (undissected) FISH, 1000 uL of freshly prepared fixation buffer (1xPBS; 3.7% formaldehyde) was added to up to 100 uL of live worms collected in an eppendorf tube. Samples were rotated at RT for 45 minutes, spun down at 3000xg in a table top centrifuge, and washed twice with 1000 uL 1xPBS. Following removal of the supernatant, 1000 uL of 75% ethanol was added to each sample, and stored at 4° C for at least 4 hours. Following ethanol permeabilization, samples were then washed once with 1000 uL of freshly prepared Stellaris® Buffer A Mixture (10% deionized formamide; 20% Stellaris® RNA FISH Wash Buffer A (Biosearch Technologies; Cat #: SMF-WA1-60); 70% RNase-free water). Following removal of the supernatant, 100 uL of freshly prepared Hybe Buffer Mixture was added (for two color *in situs*: 85.5 uL of Stellaris® RNA FISH Hybridization Buffer (Biosearch Technologies; Cat #: SMF-HB1-10); 9.5 uL deionized formamide; 2.5 uL of 5 uM probe #1 suspended in TE; 2.5 uL of 5 uM probe #2 suspended in TE) and samples were incubated at 37° C overnight. Following incubation, 1000 uL of freshly prepared Stellaris® Buffer A Mixture was added to the samples and incubated at 37° C for 30 minutes. Buffer was then removed and 1000 uL of Stellaris® Buffer A Mixture with 5 ng/mL DAPI was added, and samples were again incubated at 37° C for 30 minutes. Buffer was then removed, and samples were subsequently incubated with Stellaris® RNA FISH Wash Buffer B (Biosearch Technologies; Cat #: SMF-WB1-20) as prepared through manufacturers protocol for 5 minutes at RT. Stellaris® RNA FISH Wash Buffer B was removed and samples were resuspended in Vectashield Antifade Mounting Medium with DAPI (VWR; Cat #: H-1200-10). Samples were then placed on a slide and sealed with a coverslip and nail polish.

For FISH on dissected germlines, worms were first placed on poly-L-lysine coated slides in a solution of M9 with 10 mM levamisole to induce paralysis. Germlines were dissected with needles, and coverslips were subsequently placed over slides. Slides were then immediately frozen on dry ice, and freeze-cracked, and placed immediately into -20° C methanol for fixation. Samples were washed three times in PBS+0.1%Tween20 and fixed in 4% PFA for one hour at room temperature. Samples were then washed on the slide with 500 uL of freshly prepared Stellaris® Buffer A Mixture, followed by a brief incubation with 100 uL of Stellaris® Buffer A Mixture placed directly on the slide. Excess buffer was removed and 100 uL of freshly prepared Hybe Buffer Mixture was added (for two color *in situs*: 85.5 uL of Stellaris® RNA FISH Hybridization Buffer (Biosearch Technologies; Cat #: SMF-HB1-10); 9.5 uL deionized formamide; 2.5 uL of 5 uM probe #1 suspended in TE; 2.5 uL of 5 uM probe #2 suspended in TE) onto each slide and incubated at 37° C overnight. Following incubation, the Hybe Buffer Mixture was removed and slides were first washed with 500 uL of freshly prepared Stellaris® Buffer A Mixture and then incubated with 100 uL of Stellaris® Buffer A Mixture for 30 minutes at 37° C. Excess buffer was removed and slides were washed with 500 uL of freshly prepared Stellaris® Buffer A Mixture containing 5 ng/mL DAPI and then incubated with 100 uL of 5 ng/mL DAPI containing Stellaris® Buffer A Mixture for 30 minutes at 37° C. Following this incubation, buffer was removed, and samples were subsequently washed with 500 uL of Stellaris® RNA FISH Wash Buffer B (Biosearch Technologies; Cat #: SMF-WB1-20) as prepared through manufacturers and incubated with 100 uL of Stellaris® RNA FISH Wash Buffer B protocol for 5 minutes at RT. Stellaris® RNA FISH Wash Buffer B was removed and Vectashield was added to each sample before sealing with a coverslip and nail polish. smFISH probes were designed using the Stellaris Probe Designer (v4.2) and were purchased with Quasar670 and Quasar570 dyes. Probe sequences are listed in Table S2.

### sRNA sequencing

Small RNA libraries were prepared as follows: 5 ug of extracted total RNA was treated with 5′ polyphosphatase (20 U/ug of RNA) for 30 minutes at 37 C. RNA then was purified using the Zymo RNA Clean & Concentrator Kit columns (Zymo; Cat #: R1017). 1 ug of treated RNA was then input into the Illumina TruSeq Small RNA Library Preparation Kit (Illumina; Cat #: RS-200-0012) with 11 cycles of PCR amplification. Libraries were then run on either a 6% Novex TBE gel or a 5% Criterion TBE gel and size selected according to the Illumina protocol. Purified samples were then sequenced on the Illumina HiSeq2500 at the Johns Hopkins University School of Medicine Genetic Resources Core Facility.

### High-throughput sequencing analyses

5′ Illumina adapter sequences were removed using the default settings of Cutadapt (Martin, 2011), and reads with lengths longer than 30 nts or shorter than 18 nts were discarded. Libraries were then aligned to the UCSC ce10 reference genome using HISAT2 (Kim et al., 2015). For assessing the number of reads mapping to the *mex-6* gene, the total number of reads aligning to *mex-6* were counted and normalized to the number of singly aligned reads mapping to the genome (library size) per 1 million reads (RPM).

In order to determine if the sRNAseq libraries from our different genotypes were comparable, we compared the number of miRNA mapping reads from each mutant genotype to wild-type (Fig. S9). No drastic change in miRNA mapping reads was detected, suggesting that that there is not a global change that could affect comparisons of siRNA levels between genotypes (Fig. S9). We therefore concluded that are libraries can be accurately compared.

For sRNA read coverage analysis, mapped sRNA reads across the *mex-6* gene were placed into 5-bp bins. The number of nucleotides per bin were normalized based on library size and averaged across two technical replicates. sRNAs present in the control RNAi condition (L4440 RNAi vector) were then subtracted from the RNAi condition. All scripts are available upon request.

### pUGylation assays

pUG cDNA was synthesized using the SuperScript™ III First-Strand Synthesis System (ThermoFisher; Cat # 18080051) according to the manufacturers instruction. Briefly, 1 ug of isolated total RNA was combined with 1 uL of the 2 uM pUG specific RT primer (Shukla et al., 2020; Table S1), 1 uL of 10 mM dNTP mix, and filled up to 10 uL with RNAse-free water. Mixtures were then incubated at 65 C for 5 minutes then placed on ice for at least 1 minute. 10 uL of the cDNA Synthesis Mix (2 uL of 10x RT buffer, 4 uL of 25 mM MgCl2, 2 uL of 0,1 M DTT, 1 uL of 40 U/uL RNaseOUT^TM^, 1 uL of 200 U/uL SuperScript III RT) was then added to each reaction and then incubated for 50 min at 50 C followed by 5 min at 85 C. Reactions were then chilled on ice, and 1 uL of RNase H was subsequently added. Reactions were then incubated at 37 C for 20 min. cDNA was then stored at -20 C.

1 uL of pUG cDNA was then input into a 20 uL GoTaq PCR reaction (Promega; Cat # M7123) with the first adapter specific primer (Shukla et al., 2020; OJPO398 in Table S1) and the first gene specific primer (“f1” primers in Table S1). The PCR reactions were diluted 1:100 and 1 uL of the dilution was added to a second 20 uL GoTaq PCR reaction with the second adapter specific primer (Shukla et al., 2020; OJPO399 in Table S1) and the second gene specific primer (“f2” primers in Table S1). The reactions were then run on a 1% agarose gel and imaged with a Typhoon imager.

For pUGylation assays following the 3xFLAG::ZNFX-1 IP, 8 uL of the eluted 20 uL of RNA from the IP RNA extraction was used for the pUG cDNA synthesis (representative of approximately 40% of the IPed RNA). 5 ug of RNA was used for the IP input pUG cDNA synthesis (representative of approximately 0.25% of the input RNA). RT reactions were subsequently subjected to two rounds of PCR as described above.

### Immunoprecipitation

Adult worms were collected with a filter and washed in sonication buffer (20 mM Tris-HCl pH 7.5, 200 mM NaCl, 2.5 mM MgCl2, 10% glycerol, 0.5% NP-40, 1 mM DTT) with cOmplete™, Mini, EDTA-free Protease Inhibitor Cocktail (Millipore Sigma; Cat #: 11836170001; 1 tablet/10 mLs). Worms were flash frozen in sonication buffer and stored at -80 C. For sonication, samples were thawed on ice, and SUPERase•In™ RNase Inhibitor (ThermoFisher; Cat #: AM2694) was added for a final concentration of 80 U/mL. Samples were sonicated with a Branson Digital Sonifier SFX 250 with a microtip (15s on, 45s off, 20% power, 3 min for total on time) and cleared by centrifugation at 4 C for 15 minutes at 18,400xg. Lysate concentration was found with the Pierce BCA assay (ThermoFisher; Cat # 23225). For the IP, Anti-FLAG M2 magnetic beads (Millipore Sigma; Cat #: M8823-1ML) were prepared by vigorous vortexing. 20 uL of bead slurry was washed three times in 200 uL of sonication buffer + 80 U/mL SUPERase•In™ RNase Inhibitor, and 400 uL of 500 ug/uL lysate was added to the beads. An equivalent of 1% input lysate was used for analysis of the IP by western blot (see Western blotting). An additional equivalent of 50% of input lysate was saved for RNA extraction (see RNA extraction and purification). Samples were rotated at 4 C for 2 hours. Samples were cleared with a magnetic stand, and 4.2 uL of the supernatant (∼1%) was saved for western analysis (see Western blotting). The supernatant was removed and beads were washed 5x with 500 uL of sonication buffer + 80 U/mL SUPERase•In™ RNase Inhibitor. After final wash, beads were eluted with 4 uL of 5000 ug/mL 3xFLAG pepetide (resuspendend in TBS; Millipore Sigma; Cat #: F4799-4MG) + 96 uL of sonication buffer + 80 U/mL SUPERase•In™ RNase Inhibitor for 30 minutes at 4 C (final FLAG peptide concentration of 200 ug/mL). Beads were placed on a magnetic stand, and 1 uL of the elution was saved for western analysis (see Western blotting). Trizol was added to the rest of the elution/bead solution for RNA extraction (see RNA extraction and purification).

### Western blotting

For western blotting, DTT and Tris-Glyc SDS 2x sample buffer (ThermoFisher; Cat #: LC2676) were added to samples for a final concentration of 200 mM DTT and 1x Tris-Glyc SDS sample buffer. Samples were then flash frozen and stored at -80 C. Samples were then thawed and heated at 95 C for 10 minutes. Samples were run in Novex™ Tris-Glycine SDS Running Buffer (ThermoFisher; Cat #: LC2675) on a Novex™ WedgeWell™ 6%, Tris-Glycine, 1.0 mm, Mini Protein Gel, 12-well gel (ThermoFisher; Cat #: XP00062BOX) with a Spectra™ Multicolor High Range Protein Ladder (ThermoFisher; Cat #: 26625). Samples were transferred to an Immobilon-P PVDF Membrane (Sigma-Aldrich; Cat #: IPVH) and blocked in PBS/0.1%Tween20/5% Blotting-Grade Blocker (BioRad; Cat #: 1706404) for 30 minutes. The membrane was then incubated with anti-FLAG M2 primary antibody (1:500 dilution; Millipore Sigma; Cat #: MF1804) in PBS/0.1%Tween20/5% Blotting-Grade Blocker overnight. The membrane was washed three times for 5-10 minutes in PBS/0.1%Tween20 and incubated for 30 minutes with the Goat Anti-Mouse IgG1 HRP-conjugated secondary antibody (1:2500 dilution; JacksonImmuno; Cat #: 115-035-205) in PBS/0.1%Tween20/5% Blotting-Grade Blocker at RT. Following the secondary incubation, the membrane was washed thrice more in PBS/0.1%Tween20 and visualized with HyGLO Quick Spray Chemiluminescent HRP Antibody Detection Reagent (Denville Scientific Inc; Cat #: E2400) and the KwikQuantTM Imager (Kindle Biosciences, LLC; Cat#: D1001).

### RT-qPCR Analysis

500 ng of isolated total RNA was used as input into a 10 uL reaction of the SuperScript™ VILO™ cDNA Synthesis Kit (ThermoFisher; Cat #: 11754050), and samples were incubated according to the manufactures instructions. 3 uL of a 1:20 dilution of the cDNA was used as input into each 10 uL qPCR reaction using the SsoAdvanced™ Universal SYBR® Green Supermix (BioRad; Cat #: 1725271). *mex-6*, *puf-5*, and *oma-1* gene-specific primers were used in each reaction (final primer concentration of 250 nM for each primer). Parallel *tbb-2* qPCR reactions were run for each sample for normalization. Reactions were run on a QuantStudio^TM^ 6 Flex Real-Time PCR System (ThermoFisher; Cat #: 4485691). Fold change calculations were done based on the ΔΔCt method. Briefly, mean *tbb-2* Ct values were subtracted from the respective *mex-6*, *puf-5*, and *oma-1* Ct values (ΔCt). Average ΔCt values from the control condition of each genotype were then subtracted from the control and gene-specific RNAi condition of the same genotype (ΔΔCt). Fold change with respective to the control condition was calculated using 2^(-ΔΔCt). All qPCR primers are listed in Table S1.

### Microscopy

Fluorescence confocal microscopy was performed using two microscopes: 1) an inverted Zeiss Axio Observer with CSU-W1 Sora spinning disk scan head (Yokogawa), 1X/2.8x relay lens (Yokogawa), fast piezo z-drive (Applied Scientific Instrumentation), a iXon Life 888 EMCCD camera (Andor), and a 405/488/561/637nm solid-state laser (Coherent) with a 405/488/561/640 transmitting dichroic (Semrock) and 624-40/692-40/525-30/445-45nm bandpass filter (Semrock) respectively. Slidebook v6.0 software (Intelligent Imaging Innovations) was used for image capture, and a 63X-1.4NA objective (Zeiss) was used; 2) an inverted ZEISS LSM 880-AiryScan (Carl Zeiss) equipped with a 63X objective. ZEN imaging software (Carl Zeiss) was used for image capture, and images were subsequently processed by the ZEN Airyscan Processing method.

### Image analysis and quantification

Images were processed in Fiji (https://imagej.net/software/fiji/downloads). For *in situ* quantification of cytoplasmic RNA level, images containing both the RNA of interest (*mex-6*) and a control RNA (*puf-5*) were processed. Regions of interest (ROIs) were drawn in single Z planes corresponding to the indicated germline area, and the ROI mean intensity was calculated for both the *mex-6* and *puf-5* channels. Background mean intensity values were measured in adjacent soma tissues for both channels and subtracted from the measured values in the germline. The background-subtracted mean *mex-6* germline measurement was then normalized to the background-subtracted mean *puf-5* germline measurement, and values were plotted. Five individual worms were used for each condition.

For *in situ* quantification of pachytene nuclear signal, maximum projections were taken from half of the *C. elegans* germline and individual ROIs were drawn around 10 rows of pachytene nuclei starting in the center of the *mex-6* expression region. The maximum, mean, and median value for each ROI was measure for each channel. The median *mex-6* value for each nuclei was subtracted from its respective *mex-6* maximum value. The *mex-6* maximum value was then normalized by dividing it by the mean *puf-5* value measured for the respective ROI, and values were plotted. Three individual worms were used for each condition.

For *in situ* quantification of granule signal, ROIs for individual granules were drawn by masking in FIJI, and the mean *mex-6* and *puf-5* values was measured for each granule. Background mean intensity values were measured in adjacent soma tissues for both channels and subtracted from the measured values in the germline. The background-subtracted mean *mex-6* germline measurement was then normalized to the background-subtracted mean *puf-5* germline measurement, and values were plotted. Five individual worms were used for each condition.

---

**Figure S1. Related to Figure 1.**
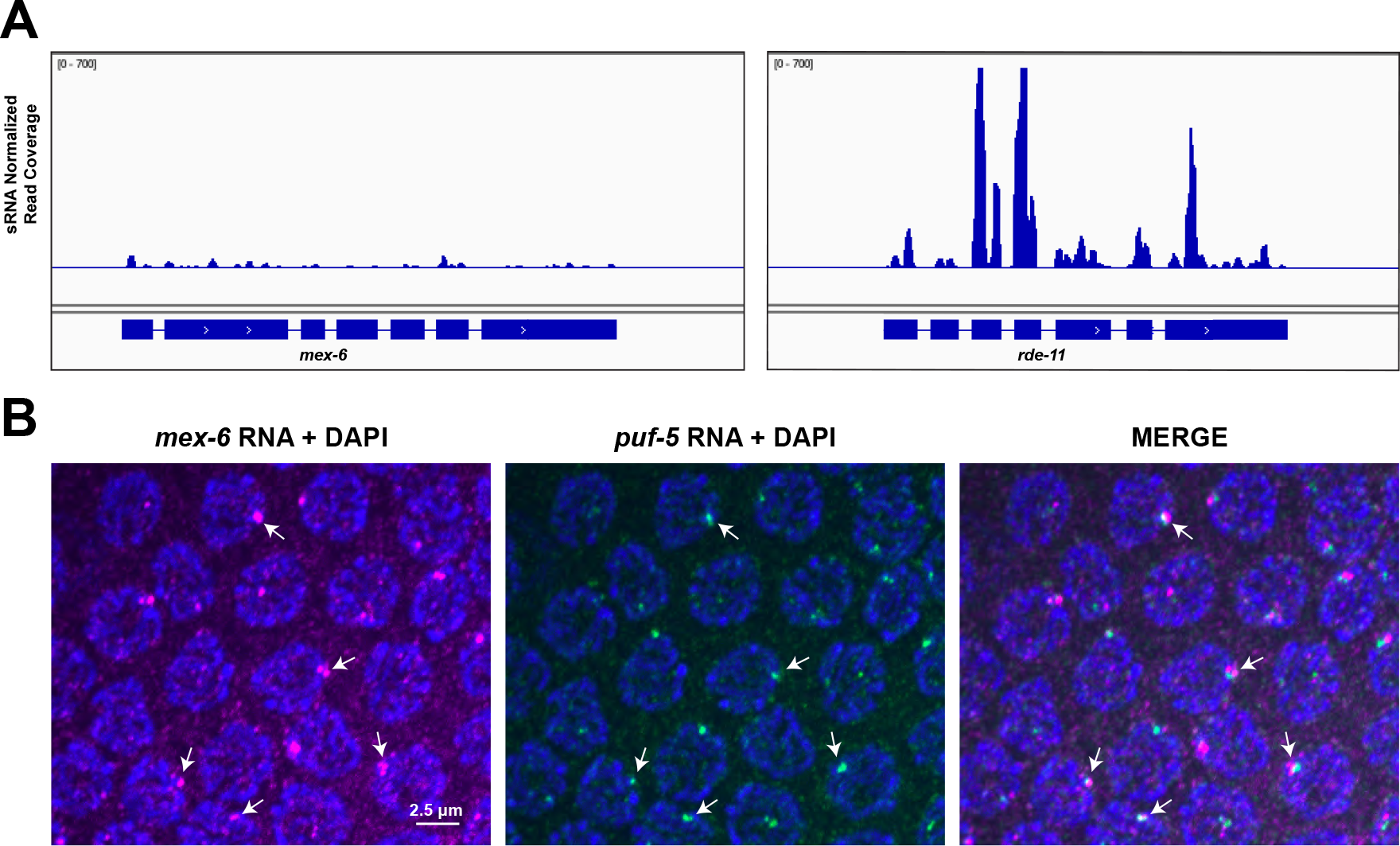
A) IGV genome browser views of sRNAseq reads across the minimally sRNA-targeted *mex-6* locus (left) and the highly sRNA-targeted *rde-11* locus (right). sRNAseq data are from wild-type adult hermaphrodites. B) Maximum projection photomicrographs of pachytene nuclei showing DNA (blue, stained with DAPI), *mex-6* RNA (magenta), and *puf-5* RNA (green). Arrows point to examples where the *mex-6* and *puf-5* RNA signals are adjacent. The *mex-6* and *puf-5* loci are located on Chromosome II less than 1 centimorgan apart.

**Figure S2. Related to Figure 2.**
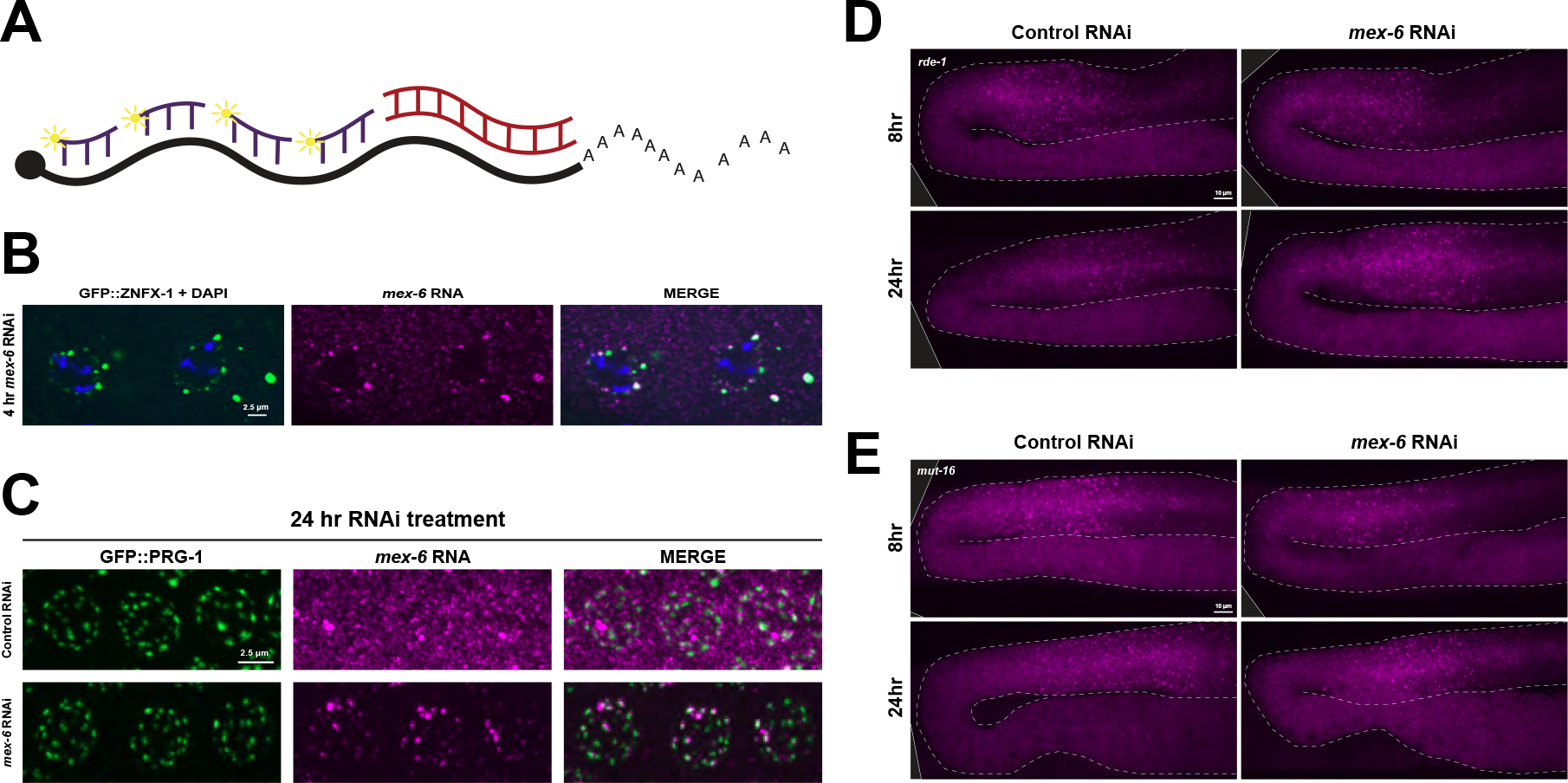
A) Schematic representation of the *mex-6* mRNA showing the smFISH probes (purple) and the dsRNA trigger (red). The dsRNA trigger targets the 3’ most 600 nts of the transcript. B) Photomicrographs of oocytes showing the Z granule marker GFP::ZNFX-1 (green), DNA (blue, stained with DAPI), and *mex-6* RNA (magenta) following 4 hours of *mex-6* RNAi treatment. C) Maximum projection photomicrographs of pachytene nuclei showing the germ granule marker GFP::PRG-1 (green) and *mex-6* RNA (magenta) following 24 hours of either control (top) or *mex-6* (bottom) RNAi treatment. D) Maximum projection photomicrographs of *rde-1* mutant germlines showing *mex-6* RNA (magenta) in either control (left) or *mex-6* (right) RNAi conditions at 8 (top) and 24 (bottom) hours of RNAi treatment. E) Maximum projection photomicrographs of *mut-16* mutant germlines showing *mex-6* RNA (magenta) in either control (left) or *mex-6* (right) RNAi conditions at 8 (top) and 24 (bottom) hours of RNAi treatment.

**Figure S3. Related to Figure 3.**
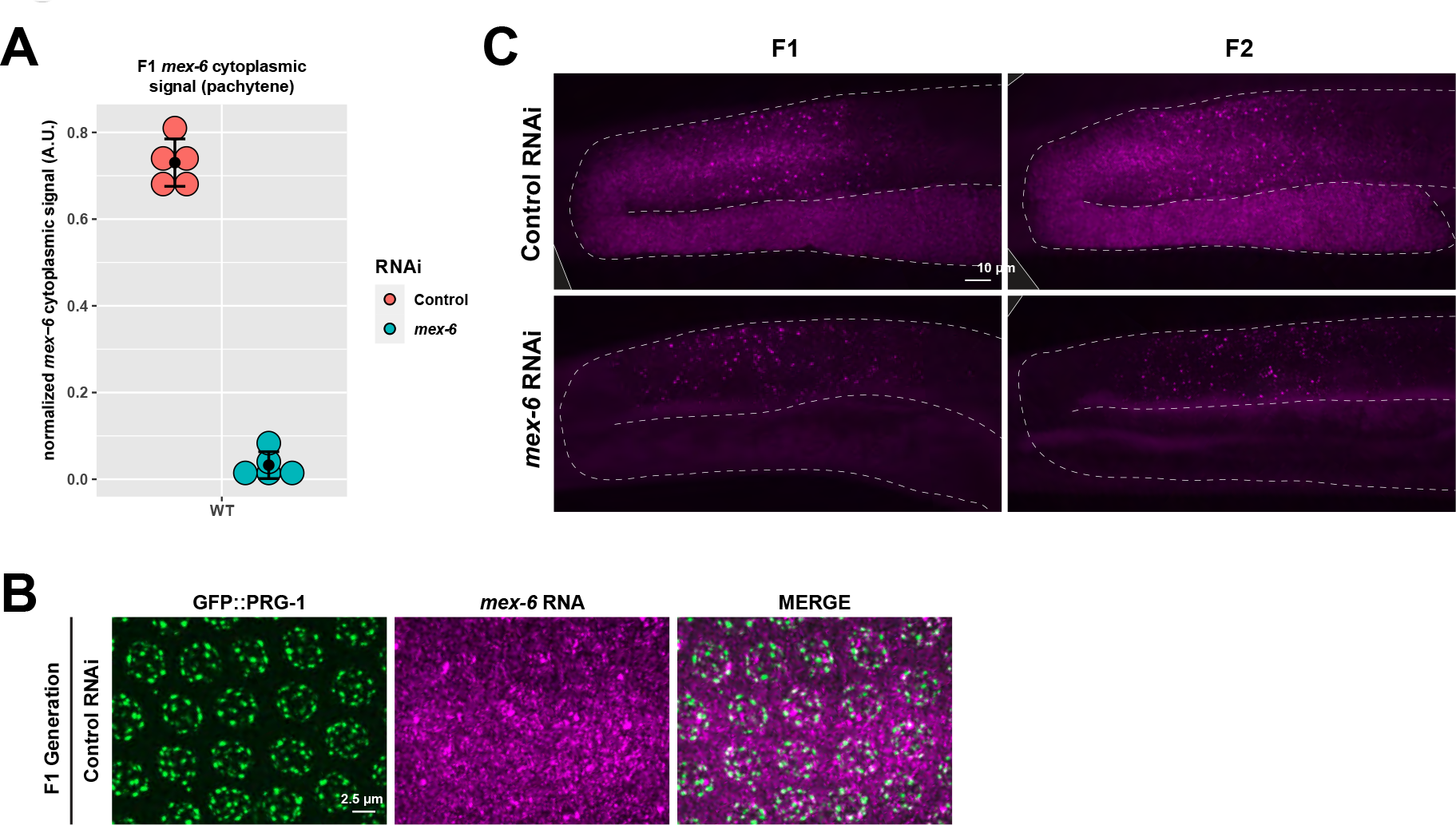
A) Graph comparing the mean *mex-6* RNA FISH signal from the pachytene rachis in wild-type F1 progeny of P0s administered either control (red) or *mex-6* (blue) RNAi. Each dot represents a single worm. Central black dot and error bars represent the mean and standard deviation, respectively. Values (arbitrary units) were normalized to *puf-5* RNA FISH signals visualized in same region (Methods) B) Maximum projection photomicrographs of pachytene nuclei showing the germ granule marker GFP::PRG-1 (green) in F1 progeny derived from animals exposed to control RNAi (*mex-6* RNAi condition for experiment shown in Fig. 3B). C) Maximum projection photomicrographs comparing germlines from F1 and F2 progeny derived from animals exposed o *mex-6* or control RNAi conditions.

**Figure S4. Related to Figure 4.**
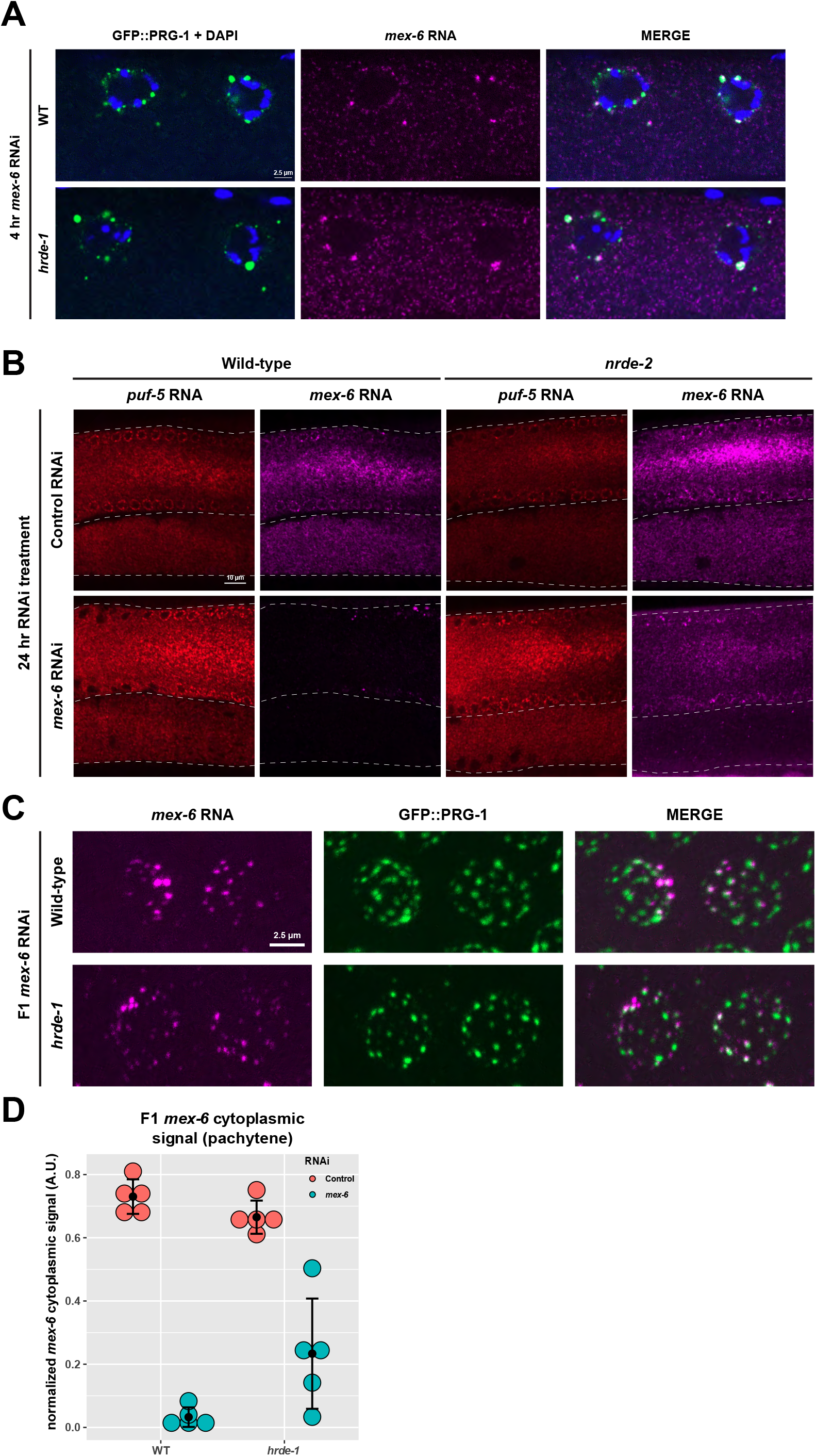
A) Photomicrographs of P0 wild-type (top) and *hrde-1*mutant (bottom) oocytes showing the germ granule marker GFP::PRG-1 (green), DNA (blue, stained with DAPI), and *mex-6* RNA (magenta) following 4 hours of *mex-6* RNAi treatment B) Single z-plane photomicrographs of P0 wild-type and *nrde-2* mutant germlines (showing both the pachytene region and oocytes) at 24 hours after control or *mex-6* RNAi administration. *mex-6* RNA FISH is shown in magenta. *puf-5* RNA FISH is shown in red (used as an in situ control). C) Maximum projection photomicrographs of F1 wild-type or *hrde-1* mutant pachytene nuclei showing the germ granule marker GFP::PRG-1 (green) and *mex-6* RNA (magenta) following administration of either control or *mex-6* RNAi in the P0 generation. D) Graph comparing the mean *mex-6* RNA FISH signal from the pachytene rachis in wild-type and *hrde-1* mutant F1 progeny of P0s administered either control (red) or *mex-6* (blue) RNAi. Each dot represents a single worm. Central black dot and error bars represent the mean and standard deviation, respectively. Values (arbitrary units) were normalized to *puf-5* RNA FISH signals visualized in same nuclei (Methods). WT values are the same as shown in Fig. S3A.

**Figure S5. Related to Figure 5.**
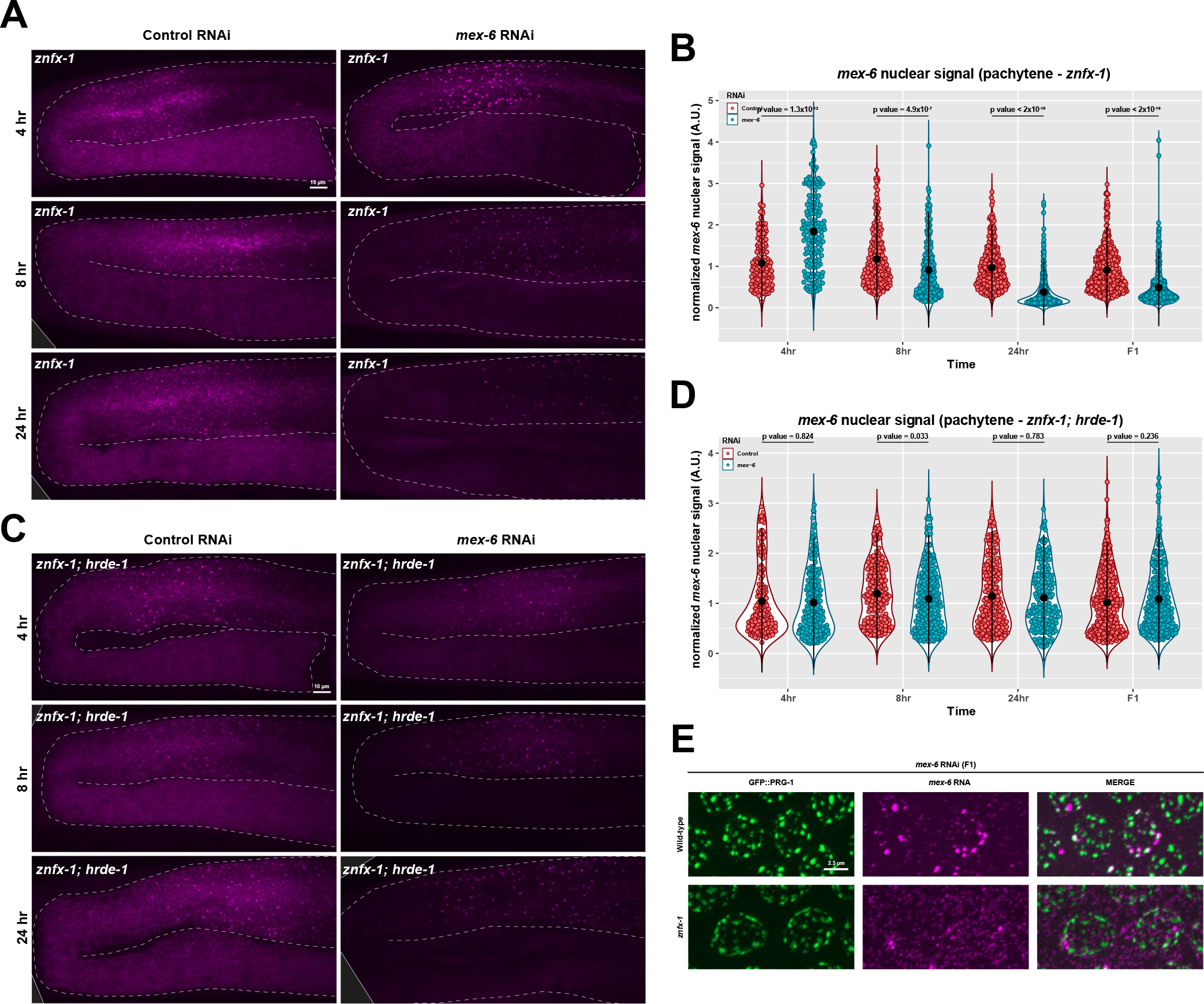
A) Maximum projection photomicrographs of *znfx-1* mutant germlines showing *mex-6* RNA at the indicated timepoints following either control or *mex-6* RNAi. B) Graph comparing the maximum *mex-6* nuclear FISH signal in control (red) vs *mex-6* (blue) RNAi at the indicated time points following RNAi in znfx-1 mutant animals. Each dot of the violin plot represents one nucleus. Nuclei in three worms were quantified for each time point and condition. Values (arbitrary units) were normalized to *puf-5* RNA FISH signals visualized in same nuclei (Methods). Central black dot and error bars represent the mean and standard deviation, respectively. P values were calculated using an unpaired Wilcoxon test. C) Maximum projection photomicrographs of *znfx-1; hrde-1* mutant germlines showing *mex-6* RNA at the indicated timepoints following either control or *mex-6* RNAi. D) Graph comparing the maximum *mex-6* nuclear FISH signal in control (red) vs *mex-6* (blue) RNAi at the indicated time points following RNAi in *znfx-1; hrde-1* mutant. Each dot of the violin plot represents one nucleus. Nuclei in three worms were quantified for each time point and condition. Values (arbitrary units) were normalized to *puf-5* RNA FISH signals visualized in same nuclei (Methods). Central black dot and error bars represent the mean and standard deviation, respectively. P values were calculated using an unpaired Wilcoxon test. E) Maximum projection photomicrographs of F1 wild-type and *znfx-1* mutant pachytene nuclei showing the germ granule marker GFP::PRG-1 (green) and *mex-6* RNA (magenta) following administration of *mex-6* RNAi in the P0 generation.

**Figure S6. Related to Figure 6.**
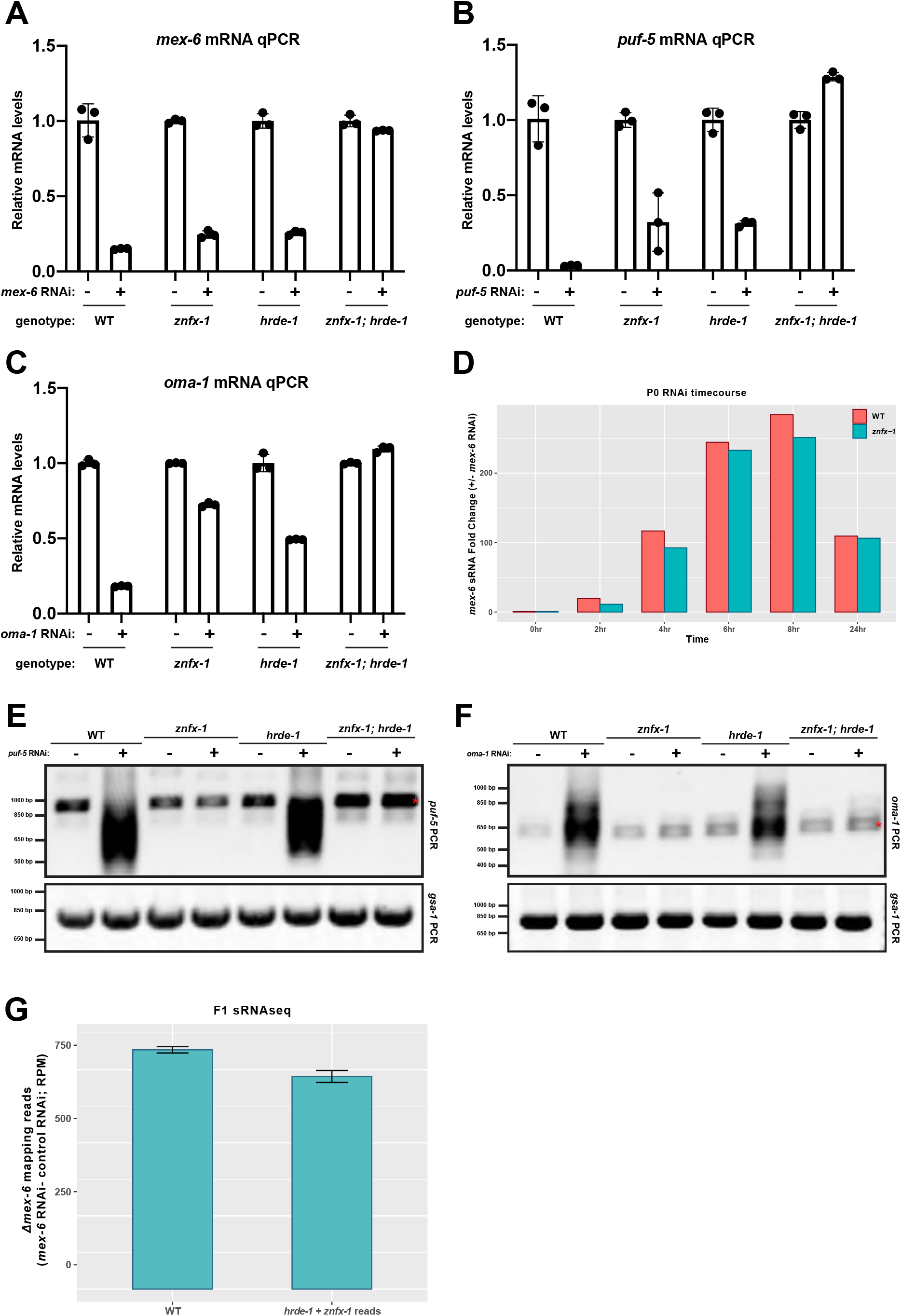
A-C) *mex-6* (A), *puf-5* (B), and *oma-1* (C) RT-qPCR from RNA derived from wild-type, *znfx-1, hrde-1,* and *znfx-1; hrde-1* F1 progeny of P0 worms administered either control, *mex-6* (A), *puf-5* (B), or *oma-1* (C) RNAi. Respective RT-qPCR Ct values are normalized to *tbb-2* RT-qPCR Ct values. The control RNAi condition for each genotype was normalized to 1 and the respective RNAi condition was compared to that (see Methods). D) Bar graph depicting the fold increase in sRNAseq reads mapping to the *mex-6* transcript in wild-type and *znfx-1* mutant P0s at the indicated timepoints following either control (red) or *mex-6* (blue) RNAi. E-F) *puf-5* (E) and *oma-1* (F) specific PCRs of pUG cDNA generated from wild-type, *znfx-1, hrde-1,* and *znfx-1; hrde-1* mutant F1s following either control (“-“) or *puf-5/oma-1* (“+”; Figure S6E/S6F, respectively) RNAi in the P0 (top panel). *gsa-1* was used as a pUG RT-PCR control (bottom panels). G) Graph comparing *mex-6* sRNA reads induced by RNAi in WT compared to the sum of the sRNA reads induced by RNAi in the *hrde-1* and *znfx-1* single mutants. See Methods for calculations.

**Figure S7. Related to Figure 7.**
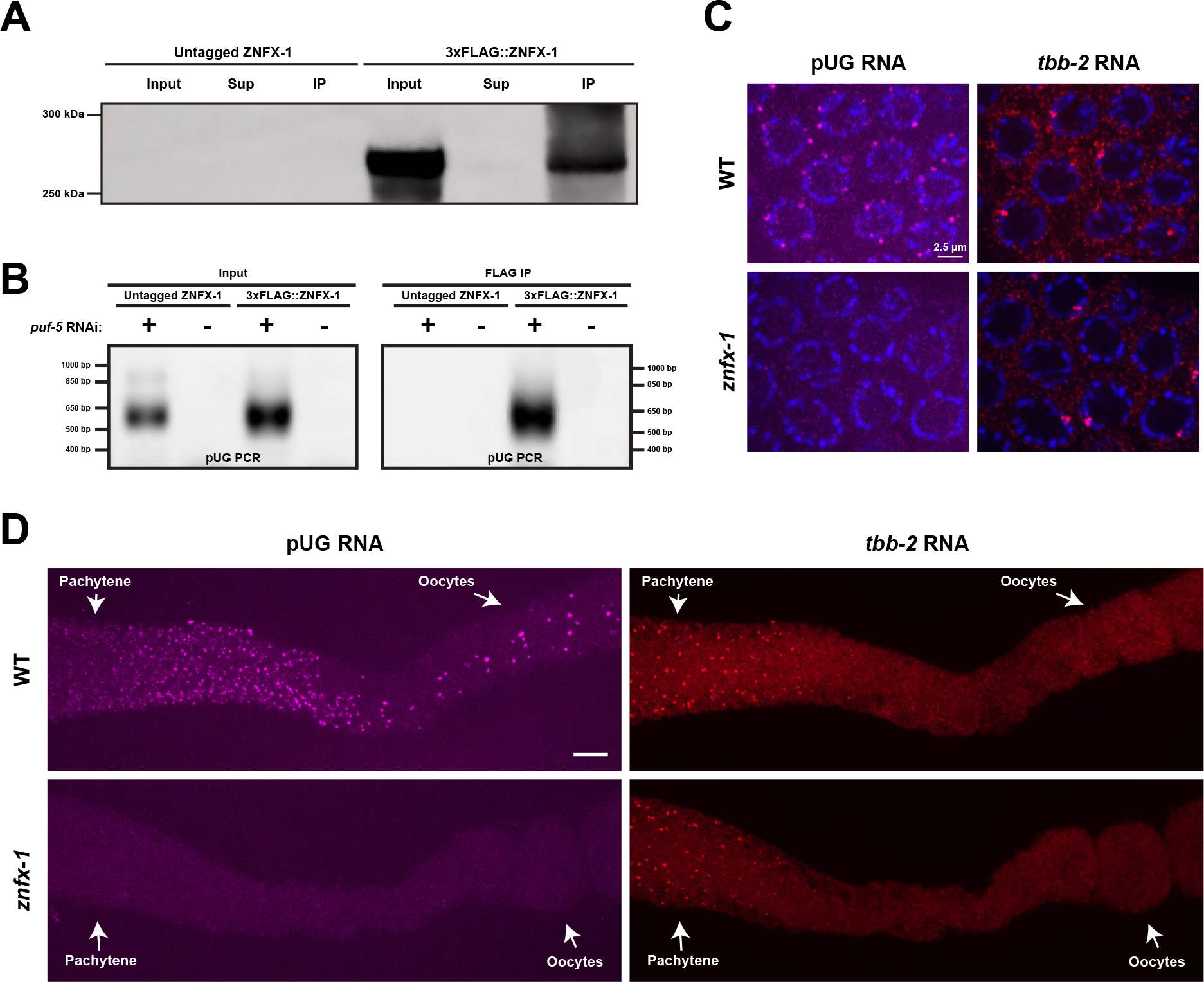
A) anti-FLAG western blot of FLAG immunoprecipitation experiments from untagged and 3xFLAG-tagged ZNFX-1 lysis. “Input” represents 1% of the input sample prior to immunoprecipitation; “Sup” represents 1% of the supernatant following immunoprecipitation; “IP” represents 1% of the immunoprecipitation sample following FLAG elution. B) *puf-5*-specific PCRs of pUG cDNA generated from input (left panel) or FLAG immunoprecipitations (right panel) of untagged or 3xFLAG-tagged ZNFX-1 lysis. Lysis was obtained from adult worms grown for 12 hours on either *puf-5* (“+”) or *mex-6* (“-”) RNAi. PCRs were generated from the same pUG cDNA used in Fig 7A. C) Photomicrographs of pachytene nuclei from dissected wild-type and *znfx-1* mutant germlines showing pUG RNA FISH (magenta), DNA (blue, stained with DAPI), and *tbb-2* RNA (red; used as an in situ control). Same images as in Fig. 7B, but equally contrasted. D) Maximum projection photomicrographs of dissected wild-type and *znfx-1* mutant germlines showing pUG RNA FISH (magenta) and *tbb-2* RNA (red, used as an in situ control). The pachytene region and oocytes are indicated with arrows.

**Figure S8. Related to Discussion.**
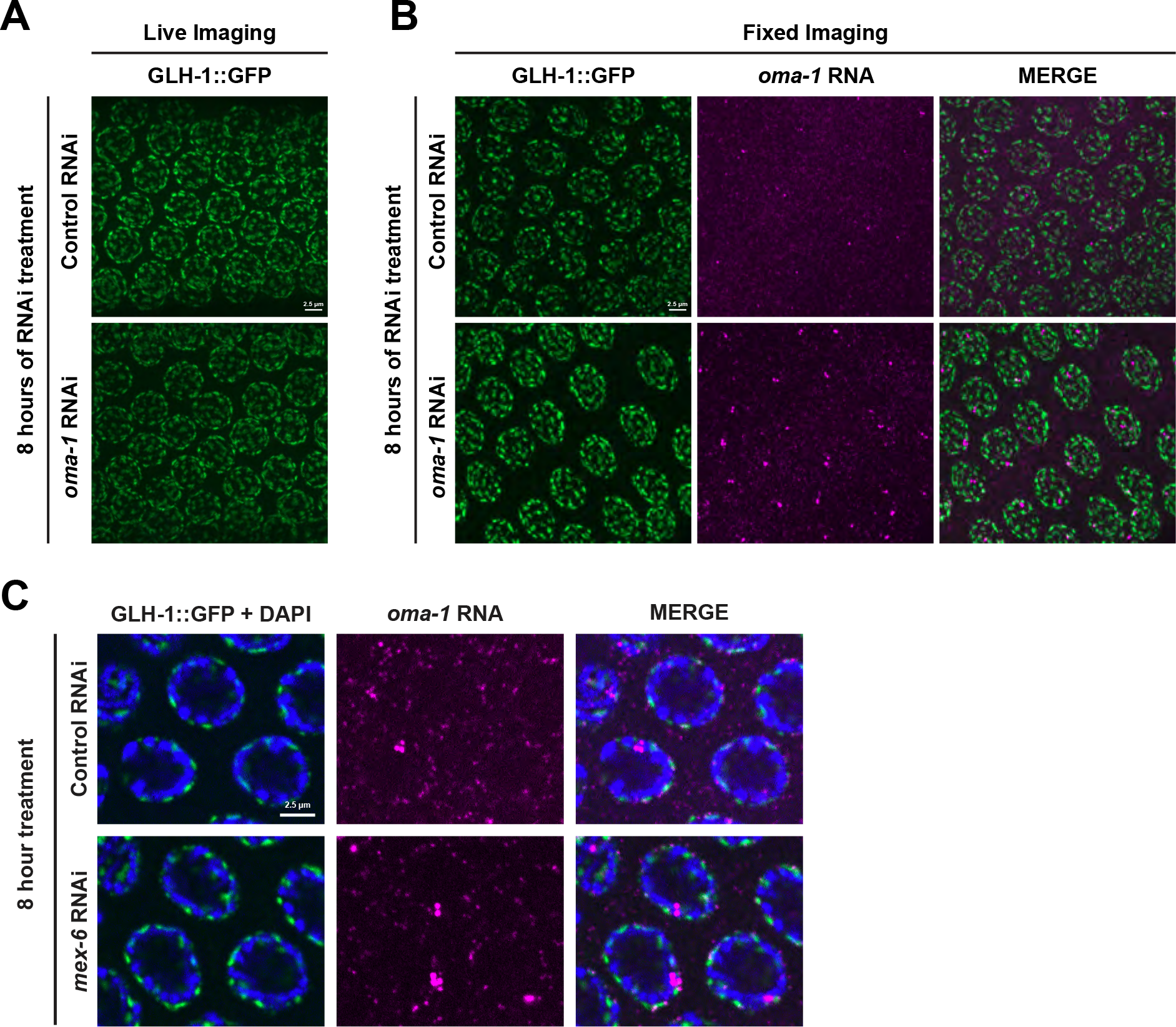
A) Maximum projection photomicrographs of live-imaged pachytene nuclei showing GLH-1::GFP after 8 hours of either control or *oma-1* RNAi treatment. B) Maximum projection photomicrographs of fixed-imaged pachytene nuclei showing GLH-1::GFP (green) and *oma-1* RNA (magenta) after 8 hours of either control or *oma-1* RNAi treatment. Note that the *oma-1* RNA foci do NOT overlap with GLH-1::GFP foci. C) Single Z-plane photomicrographs of fixed-imaged pachytene nuclei showing GLH-1::GFP (green), DNA (blue, stained with DAPI) and *oma-1* RNA (magenta) after 8 hours of either control or *oma-1* RNAi treatment. The *oma-1* RNA foci overlap with DAPI and not with with GLH-1::GFP foci.

**Figure S9. Related to Methods.**
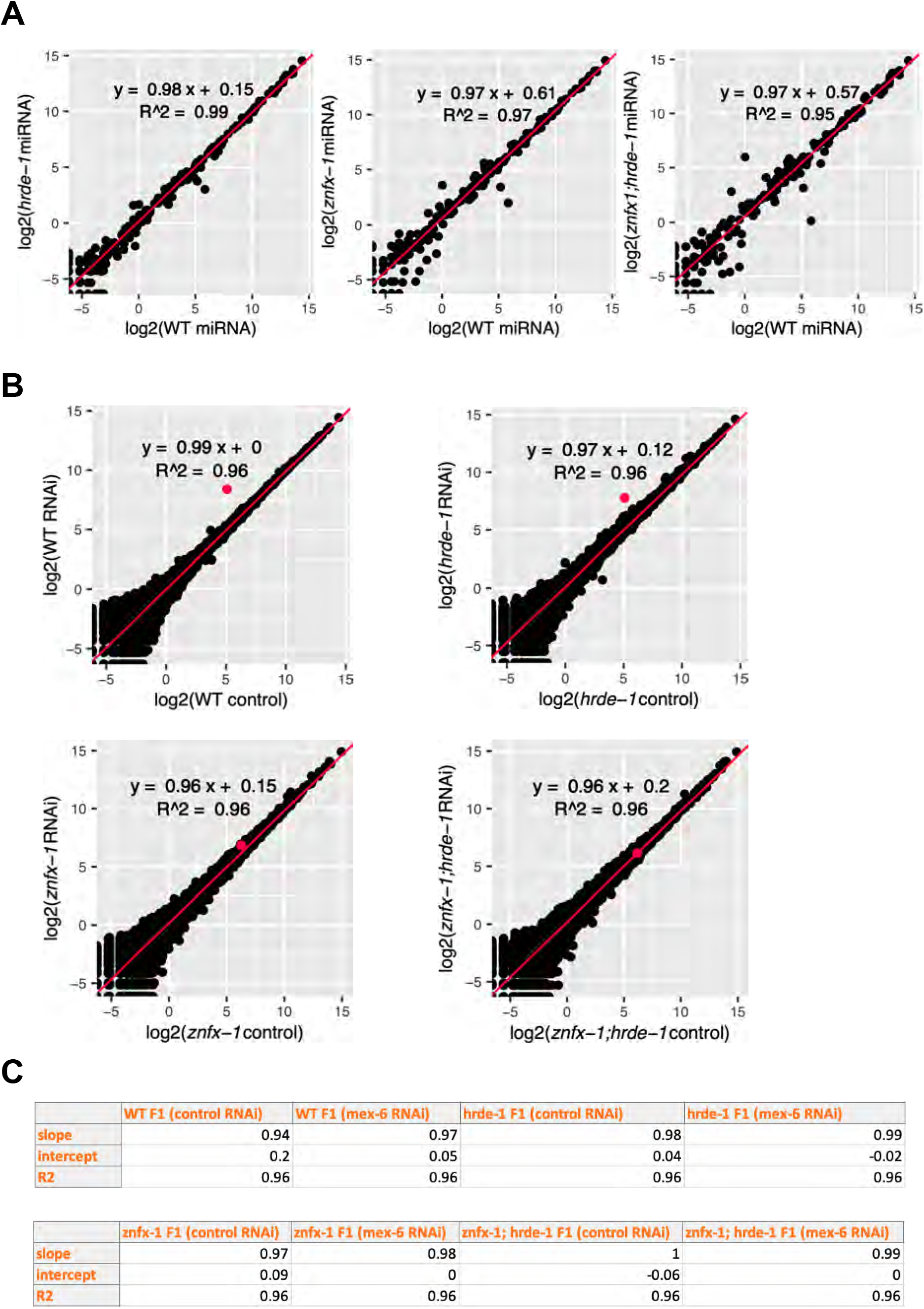
A) Scatter plots comparing miRNA RPMs in wild-type (X-axis) and mutants (Y-axis) as indicated under control RNAi conditions. Linear regression is used to fit the data. miRNA counts in each condition are averaged across two replicates. B) Scatter plots comparing sRNA RPMs in the F1 progeny of control and mex-6 RNAi fed P0 worms for the indicated genotypes. Each dot corresponds to a locus in the *C. elegans*genome. The red dot corresponds to the *mex-6* locus. Linear regression is calculated without *mex-6* sRNA counts. sRNA counts in each condition were averaged across two replicates. C) Linear regression statistics modeling the relationship between the two sRNAseq replicates for each genotype in both control and *mex-6* RNAi conditions.

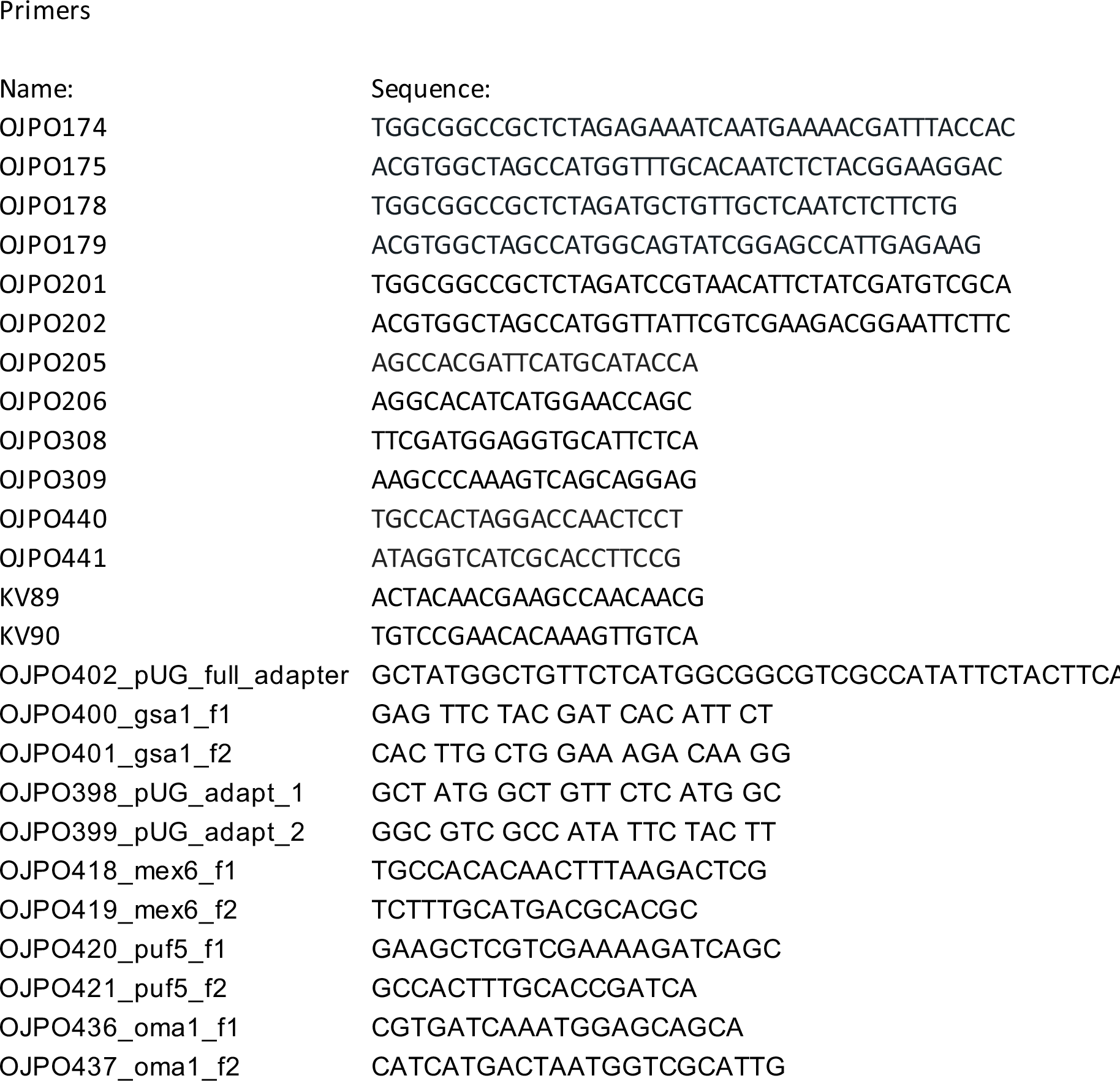
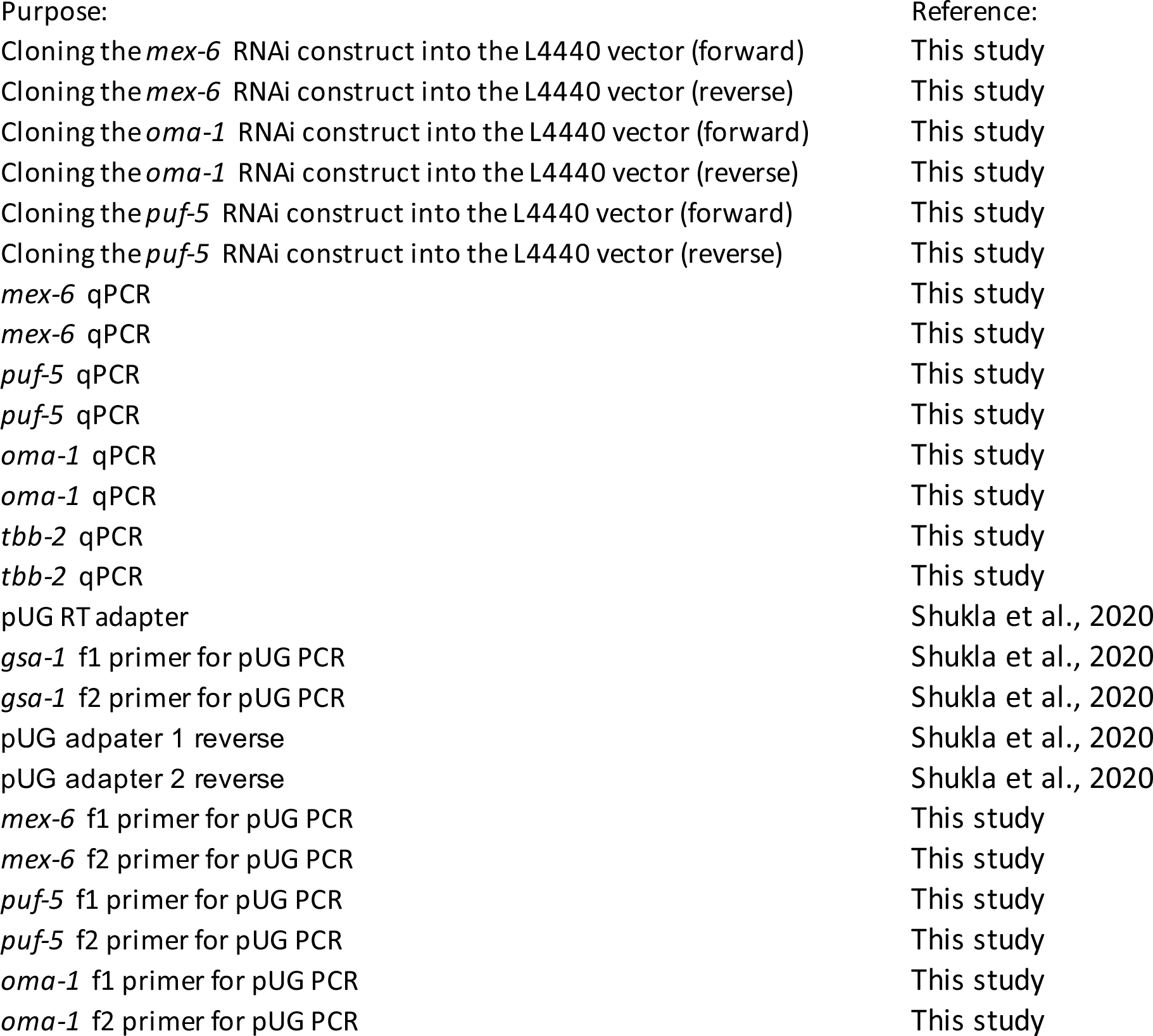
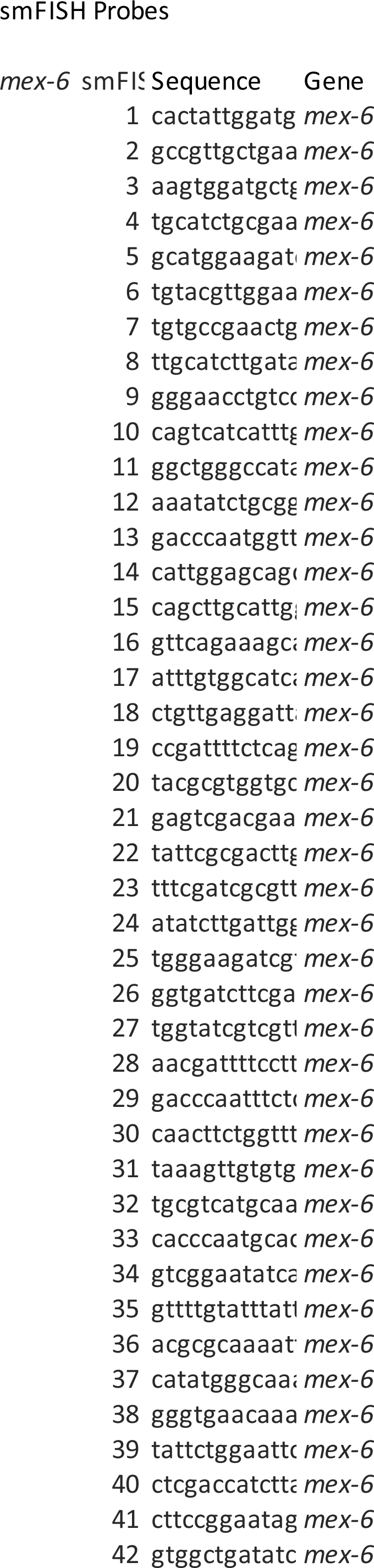
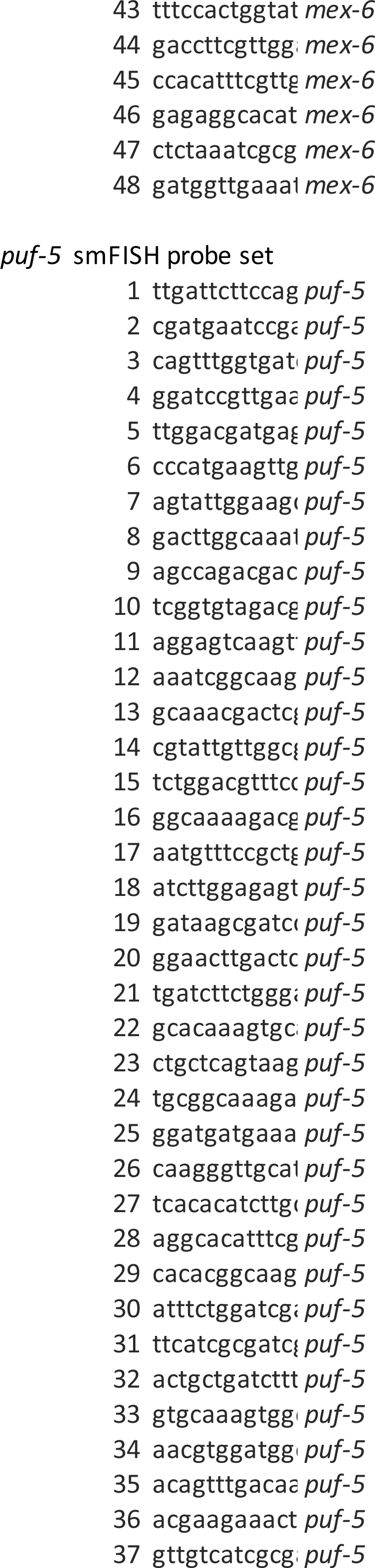
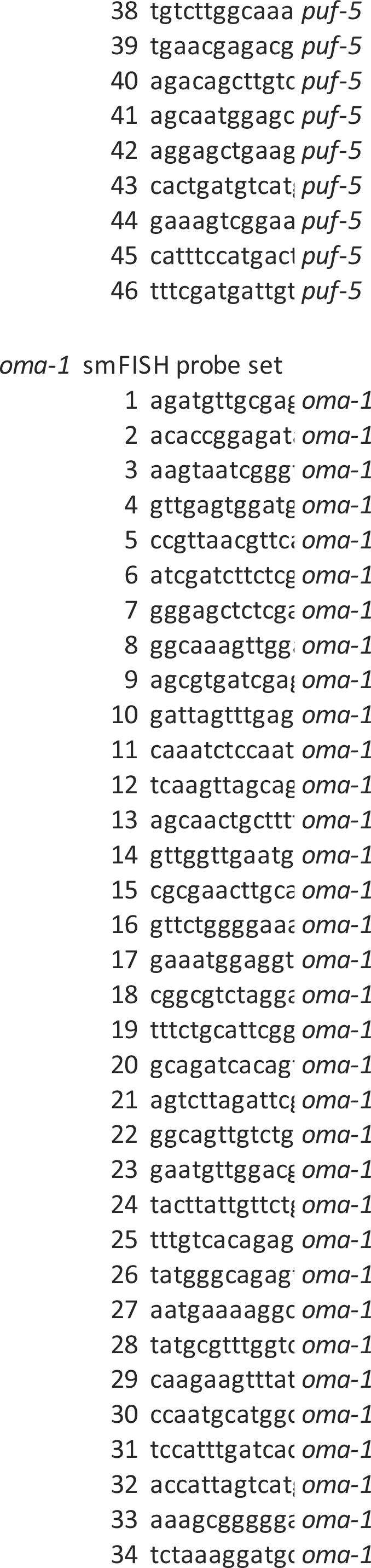
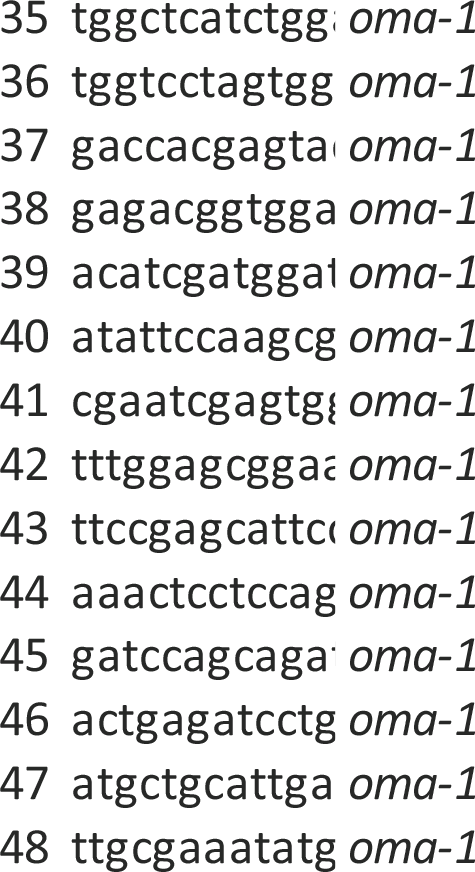

